# Tunable population diversity through a noise-driven differentiation switch

**DOI:** 10.1101/2025.09.20.677552

**Authors:** Jialong Jiang, Sisi Chen, Jong H. Park, Tiffany Tsou, Vickie Yang, Paul Rivaud, Matt Thomson

## Abstract

During development, combinatorial signals modulate progenitor differentiation to build populations of diverse cell types. Classical deterministic fate-control circuits focus on stable commitment but do not explain how a diverse cell population is generated and tuned by signals. Here, using single-cell RNA-seq profiling of neural progenitor cells (NPCs) upon combinatorial signal inputs, we identify a candidate circuit-level mechanism for controlling cell type compositions through an annealed noise-driven switch. With 40 signaling conditions, we show that combinatorial signal inputs modulate the neuronal-glial population proportions following a simple log-linear model. Regulatory network models built by D-SPIN quantitatively capture cell state distribution shifts induced by signal combinations, and indicate a circuit model for the probabilistic fate specification by a differentiation switch with controlled transcriptional heterogeneity. Noise levels drive the system from highly plastic to committed states, allowing the relative level of signal inputs to shift population structures. The model further predicts an early bipotent state expressing lineage-specifying factors of both fates, which we verify in our single-cell profiling experiment and a mouse embryonic visual cortex development dataset. Our work identifies a design principle of noise-driven additive regulation in the logarithmic cell-fate probability space, providing a new strategy for population-level stem cell control.

## Introduction

Organogenesis depends on the coordinated generation of diverse cell types from progenitors that integrate their signaling environment into lineage choices [1, 2]. At the individual cell level, signal inputs are processed through interactions between gene regulators, which define canalized cell states and govern transitions between them [3, 4, 5, 6]. However, the expression of these regulators have been found to be heterogeneous and dynamic, predisposing progenitor cells to distinct differentiation outcomes under the same signaling cues [7, 8, 9, 10, 11]. A central challenge is to understand how combinatorial signal inputs reshape the distribution of differentiation outcomes of progenitors, and how these population-level responses emerge from the interplay between transcriptional variability and fate-control circuits.

Stem and progenitor cell culture provides a tractable system for investigating mechanisms of signal integration and cell fate modulation, allowing the signaling environment to be systematically varied [12, 13, 14]. Neural progenitor cells (NPCs) can be extracted from mouse embryos and cultured as neurospheres under growth factor supplementation, and induced to generate neurons, astrocytes, and oligodendrocytes [15, 16]. During *in vivo* brain development, numerous mechanisms contribute to shaping the relative abundance between neuronal and glial lineages, such as temporal competence switching, spatial patterning and feedback cell-cell interactions [17]. Complementary *in vitro* cell culture studies have shown that extracellular signals can bias NPC differentiation toward neuronal or glial lineages [18, 19]. The controlled environment and multilineage potential set up a suitable system to investigate how shared signals are interpreted by heterogeneous progenitors.

In this study, with single-cell RNA-seq profiling, we discover a potential mechanism of population control through combinatorial signal inputs on a heterogeneous progenitor population. We find that the relative levels of EGF/FGF2, BMP4, and Wnt can continuously tune the differentiation outcomes of NPCs from glia-dominant to neuron-dominant (Figure 1A). Among 40 combinatorial signaling conditions, the logarithmic ratio between neuronal versus glial proportions is a linear combination of individual signal actions. Using a network inference framework, D-SPIN [20], we uncover distinct signal-specific genes and gene programs that reshape the cell state distribution, such as Neurod1/6, Olig1, Hes1, and Sp9. The inferred circuit structures motivate an annealed noise-driven differentiation switch model in which signals bias two alternative fate states, while a time-dependent effective noise level controls transitions between them (Figure 1B). The model recapitulates the observed log-linear signal integration and predicts an early bipotent progenitor state expressing regulators of both fates, which we validate in both our single-cell experiment and a public mouse embryonic visual cortex development dataset [21]. Our work experimentally identifies a design principle of additive signal integration in the log-ratio space, and proposes a circuit-level mechanism for population composition control by combinatorial signal inputs.

**Figure 1.**
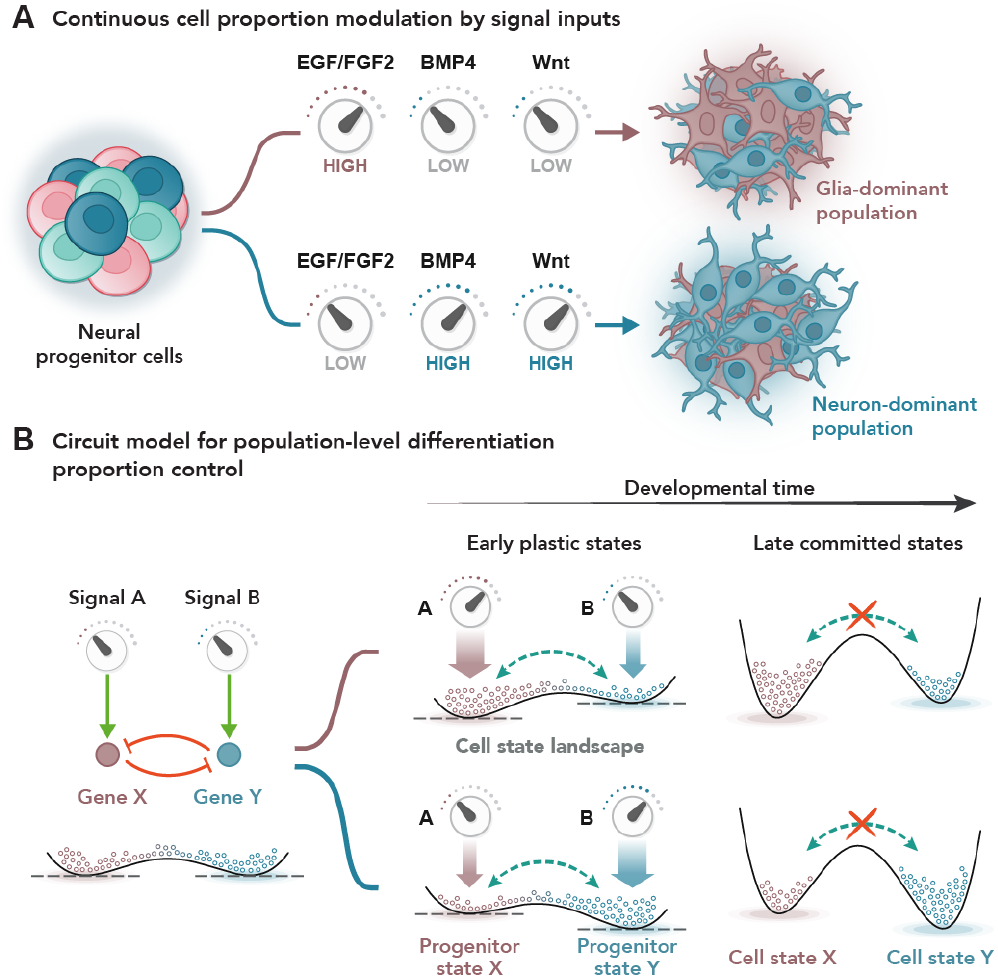
Annealed noise-driven switch generates tunable population composition with combinatorial signal inputs. (A) Schematics for continuous modulation of population composition through combinatorial inputs. Signal inputs such as EGF/FGF2, BMP4, and Wnt can tune the neuron-glia proportion of neural progenitor cell differentiation outcomes. (B) Schematics for circuit mechanism of population-level differentiation control. Two fate regulators, genes X and Y, mutually repress each other and receive signal inputs A and B. In early plastic states, high effective noise permits transitions between the two states, allowing the signal inputs to tilt the cell-state landscape and continuously tune the occupancy of progenitor states X and Y. As effective noise decreases over developmental time, transitions between states are suppressed, stabilizing the proportion between committed cell states X and Y.

## Results

### Combinatorial signals tune neuronal-glial population composition in NPC culture

To understand how a heterogeneous progenitor population responds to signal inputs, we used an NPC population isolated from E18 mouse embryo SVZ and selected a range of signaling conditions for evaluation (Figure 2A, Supplemental information). Apart from the canonical EGF/FGF2 ligands used in NPC culture, we used signaling cues of BMP4, Wnt, PDGF, and FGF9 that are known to regulate diverse functions during brain development such as proliferation (EGF/FGF2), neuronal differentiation (Wnt, PDGF), astrocytic differentiation (BMP4, FGF9), and the maintenance of quiescence (BMP4, Wnt) [19, 22, 23, 24, 25, 26]. We also included two immune signaling cytokines, GM-CSF and IFNG, as outgroup controls that are not normally active during neural development.

**Figure 2.**
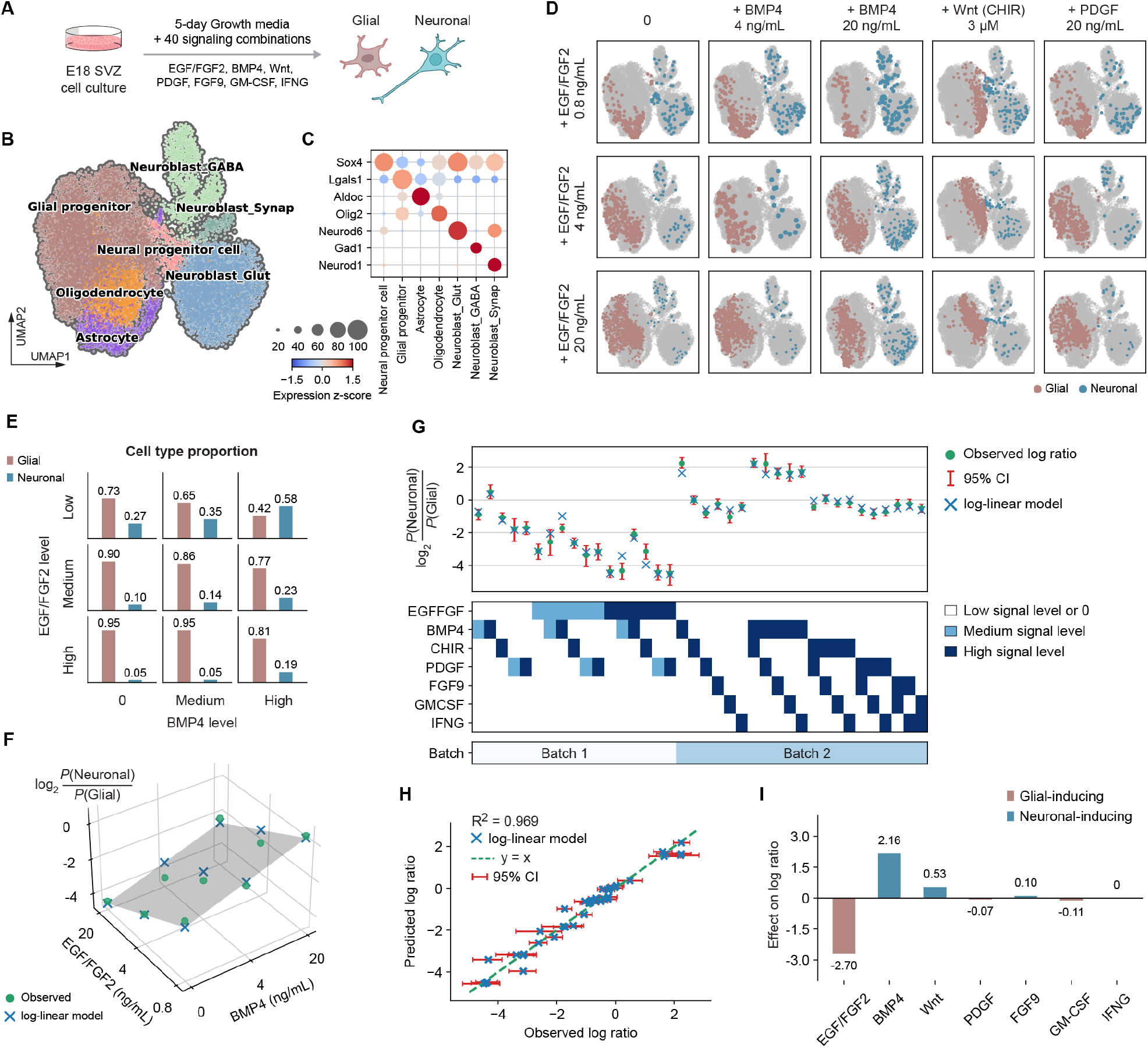
Signal combinations modulate neuronal-glial proportion following a simple log-linear model. (A) Experimental schematics showing the procedure of signal combination profiling. (B) UMAP embedding of 20k quality-controlled single cells obtained from the signal combination response experiments. (C) Bubble plots showing marker gene expression z-score per cell type. The dot sizes and colors encode the fractions of cells with detected expression and average expression levels. (D) UMAP embedding of the cell population treated with selected signals and their combinations, including different levels of EGF/FGF2 and BMP4, CHIR (Wnt activator), and PDGF. (E) Bar plots showing the proportion of neuronal cells and glial cells under different level combinations of EGF/FGF2 and BMP4. (F) The logarithm ratios of neuronal versus glial cells (log ratios) respond linearly to the combinatorial signal input of EGF/FGF2 and BMP4. The gray plane and blue crosses are predictions of a linear model, and green dots are observed log ratios. (G) The log ratios respond linearly to the signal combinations across the profiled signaling conditions. Concentrations of each signal are normalized to up to three levels: low (0), medium (0.5, optional), and high (1) as the linear model input. Confidence intervals (CIs) are computed from the bootstrap variation of each condition. A batch indicator was included as an additional regression variable to account for the systematic offset between experimental batches. (H) Scatter plot of observed log ratios versus predicted log ratios by the linear model. The log-linear model agrees well with the data across a wide range of neuronal/glial cell proportions. (I) Coefficients of the linear model on the response of cell type proportion log ratios. EGF/FGF2 induces increased glial proportions, while BMP4 and Wnt induce increased neuronal proportions.

Without signal stimuli, canonical EGF/FGF2 culture rapidly narrows the observed cell state distribution toward the glial lineage. For reference, we performed pilot experiments to profile the dynamics of cell state distributions of the SVZ population during *in vitro* cell culture and compared them with *in vivo* brain development (Figure S1A, Supplemental information). We used brain samples of three stages, E18, postnatal day 4 (P4), and adult cortex. Single-cell RNA-seq revealed a heterogeneous starting population containing neural progenitors alongside neuronal and glial cells (Figure S1B and S1C). Different from the *in vivo* diversification of specialized cell types, EGF/FGF2 culture drives a rapid and nearly complete concentration to glial states. The glial populations rise from 6.0% to 83.7% at day 3 and continue to rise to 99.5% at day 18 (Figure S1D and S1E).

Media-switching experiments confirmed that the glial states reflect a restriction in the neurogenic capacity rather than a reversible bias imposed by the current signaling environment. E18 SVZ cells cultured immediately in serum-containing differentiation media for 5 days could generate a mixed culture of 49.8% glial cells and 29.8% neuronal cells (Figure S1F and S1G). However, the cell population expanded for 5 days in EGF/FGF2 growth media remained dominated by glial cells after the subsequent treatment with differentiation media, comprising 98.2% glial cells and only 0.4% neuronal cells (Figure S1G and S1H). These experiments reveal a time-dependent transition from a heterogeneous, neurogenic-competent progenitor population to a glia-dominant population with restricted responsiveness to subsequent neuronal differentiation cues.

Informed by the time course and media-switching experiments, we exposed freshly dissociated E18 SVZ progenitor population to 40 combinatorial signaling conditions and profiled the response with single-cell RNAseq. We selected three doses of EGF/FGF2 for combination ranging from 20 ng/mL, a high concentration for standard culture conditions, down to 0.8 ng/mL, a concentration that we found was necessary to maintain cell survival. BMP4 and PDGF signals were administered at two dosages, and other signals were administered at one selected dosage (Supplemental information Table). The Wnt pathway was activated with CHIR99021 [27]. Cells were profiled with the ClickTags multiplexing platform after 5 days of exposure, similar to the setup of a parallel study (Supplemental information) [28]. Across signaling conditions, the identified cell types are highly congruent with the previous *in vivo* and *in vitro* differentiation experiments with cluster-specific expression of a similar set of marker genes (Figure 2B and 2C, Supplemental information). Cell populations fall into two major subpopulations, including glial states (glial progenitors, astrocytes, oligodendrocytes) and neuronal states (NPCs and neuroblasts of different neuron types).

Single-cell expression profiles reveal how signaling conditions shift the balance between glial and neuronal populations. Three individual signals significantly alter the abundance of neuronal and glial populations. Increasing EGF/FGF2 dosages increases the glial state abundance, while BMP4 and Wnt increase neuronal state abundance (Figure 2D). Population-level effects of other signals are less perceptible. Apart from modulating the neuronal/glial abundance, BMP4 also enriches astrocyte states in the glial population, and Wnt promotes specific glial progenitor states that occupy a distinct region on the UMAP embedding. The populations treated with signal combinations share similar signatures of both single signals. Specifically, the neuronal/glial proportions of signal combinations lie between those of individual signals. For example, the control population with minimal EGF/FGF2 has 26.9% neuronal cells; high EGF/FGF2 has 4.6%, and high BMP4 has 58.5%; the combination of high EGF/FGF2 and BMP4 produces 19.1% neuronal cells (Figure 2E).

### Combinatorial signals control neuronal-glial proportion following an log-linear model

Strikingly, the neuronal-glial proportions under combinatorial inputs can be predicted from the single-signal responses following a simple log-linear model. The logarithm ratio between neuronal proportion and glial proportion (denoted as log ratio for simplicity), log_2_ *P* (Neuronal)*/P* (Glial), falls into a three-dimensional plane when plotted against the input level of EGF/FGF2 and BMP4 (Figure 2F). We extended the analysis to the 40-condition screen collected in two experimental batches. Fitting a single parameter for each of the 7 different signals and batch effect in the log-linear model produces highly accurate predictions about the population compositions (Figure 2G). The log ratio model outperforms other candidate linear models (Figures S2A and S2B). The observed versus model-predicted log ratios align well with *R*^2^ = 0.969 (Figure 2H). The simple model contains no interactions between signals, and individual signal contributions suffice to capture lineage outcomes across a 2^6^ = 64 fold dynamic range of neuronal/glial ratios. Including signal interactions into the regression does not statistically significantly improve the prediction accuracy (Figures S2C and S2D). The accuracy of the simple log-linear model suggests that the additive impact of signaling molecules could be sufficient to explain much of the population restructuring observed across the combinatorial signaling conditions assayed in our screen.

The coefficients of the log-linear model quantify the effects of each signal on the neuronal/glial population structure modulation and enable rational design of population compositions (Figure 2I). The coefficient at the log_2_ scale translates to cell abundance fold change by computing the exponents of 2. EGF/FGF2 strongly promotes glial states by nearly 6.5-fold. BMP4 induces neuronal states by around 4.5-fold, and Wnt slightly biases neuronal states by 1.4-fold. Other signals, including PDGF, FGF9, GM-CSF, and IFNG, have little effect on the cell population structure. The formulation of a log-linear response provides a rational design strategy to achieve desired population structures through combinatorial signal inputs.

The log-linear law further indicates that the underlying regulatory mechanism allows additive signal integration for continuous cell proportion tuning. Additivity could arise if each signal linearly modulates the growth rate or survival rate of each lineage [22], or the signals may reshape the landscape of transcriptional states by shifting the relative free-energy barrier heights between neuronal and glial attractors [29]. Single-cell transcriptome profiling provides the fine-grained resolution to look beyond cell type counts and interrogate how genes and pathways respond to combinatorial signaling cues.

### Program-level network model recapitulates additive cell fate modulation effects

To understand how population-level additive signal responses are implemented at the transcriptional level, we applied the computational framework D-SPIN to construct a regulatory network model of major cellular activities and how external signals modulate them [20]. For interpretability, we first constructed a network model of gene programs, i.e., co-expressing and non-overlapping groups of genes across conditions. We identified 14 gene programs (P1-P14) by the gene program discovery pipeline of D-SPIN, followed by manual curation (Figure 3A, Figure S3A, Supplemental information). The 14 programs reflect major biological activities in the cell population, including housekeeping (P3), mitosis (P5), and lineage-specific programs of glial cells and different neuron types (P1, P12, P13). The program set also includes signal-specific programs, such as P8 and P9, that are primarily induced by Wnt and BMP4 signaling, respectively.

**Figure 3.**
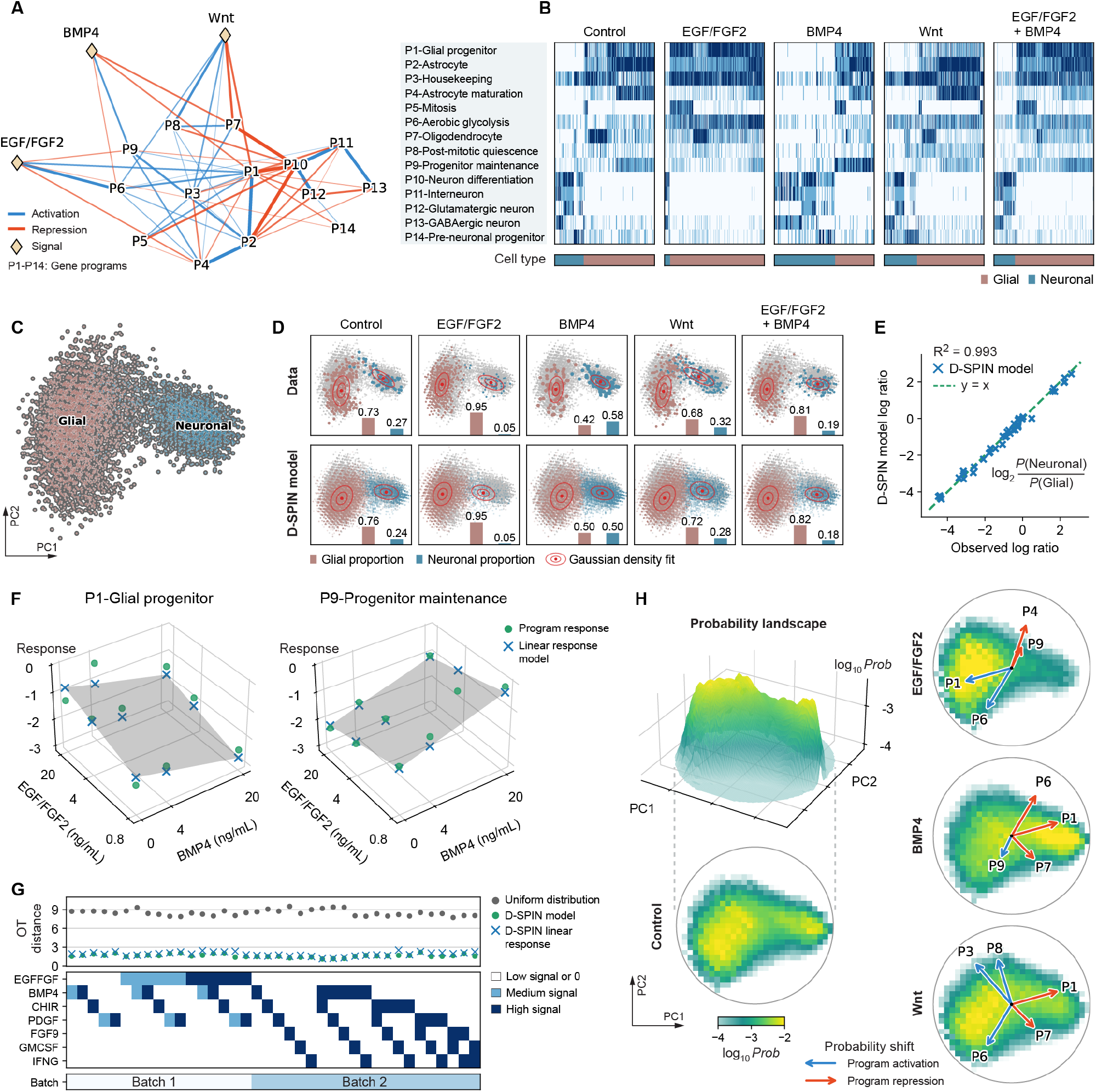
Program-level network model reconstructs cell state distributions and dissects signal responses. (A) Diagram of program-level network model inferred by D-SPIN shows interactions between 14 gene-expression programs (P1-P14, circles) and the most impactful signals, EGF/FGF2, BMP4, and Wnt (diamonds). Interactions are rendered as blue (activation) and red (repression) edges, with thickness scaled with the strength of interaction. (B) Heatmaps showing program-expression profiles of example signal and signal combination conditions. (C) PCA projections of the pooled program-level expression for the cell population treated by signal and signal combinations. (D) PCA projections of (top) experimental data and (bottom) reconstructed state distribution by the D-SPIN network model for example conditions. Inset bar plots quantify the glial and neuronal cell proportions in data and model-generated distributions. Red contours indicate Gaussian density fits to glial and neuronal cell states separately. (E) Scatter plots of observed log ratios between neuronal and glial cell proportions versus the log ratio of distributions generated by the D-SPIN model. (F) D-SPIN-inferred program-level responses are linearly modulated by the signal combination inputs, as shown in the two example programs under different EGF/FGF2 and BMP4 conditions. The gray plane and blue cross are predictions of a linear response, and green dots are inferred responses from data. (G) Scatter plots showing optimal transport (OT) distance between experimental data and cell-state distributions generated by D-SPIN, compared with the uniform distribution as a null model. Both of the DSPIN models, either with no constraints or constrained to have linear signal combination responses, have much lower OT distance (1.62, 1.88 on average) compared with the null model (8.47 on average). (H) Heatmaps of the model-generated probability landscape on the PCA projection space for (left) control and (right) distributions generated by linearly regressed responses for EGF/FGF2, BMP4, and Wnt signals. As PCA is a linear operation, the program-level responses can also be projected onto the landscape to indicate their modulation effect on probability densities, as rendered as blue (activation) and red (repression) arrows in the PCA space.

The network model inferred by D-SPIN provides an interpretable wiring diagram between internal cellular processes as well as the responses to external signal inputs (Figure 3A, Figure S3B). The network contains two subnetworks or modules, one containing glial-associated programs and the other containing neuronal-associated programs, where intra-module interactions are predominantly activating and inter-module edges are inhibitory. This architecture supports internally coordinated lineage programs while opposing simultaneous activation of the alternative fate program [30]. The three signals with significant population remodeling effects, EGF/FGF2, BMP4, and Wnt, each exhibit distinct interaction signatures with the regulatory network. Notably, these signal-to-program interactions are concentrated in the glial module, suggesting the shifts in neuronal-glial proportions are effected primarily through glial program activities.

Program-level cellular activities provide a finer-grained signature of how each signal reshapes cell states. Consistent with the inferred regulatory interaction from signals, program-expression profiles show that each signal activates distinct biological functions (Figure 3B). Compared with the control condition, EGF/FGF2 and Wnt elevate the aerobic glycolysis program (P6). The metabolic switch to glycolysis plays important roles in neural development, and also correlates with energetically demanding, proliferative progenitor states [31, 32]. BMP4 activates P9, a program relevant to progenitor maintenance containing Id1/2/3, Mdk, and Hes5, agreeing with previous studies on BMP4 supporting stem cell self-renewal [33]. Wnt also activates the housekeeping program (P3) of ribosomal and mitochondrial genes, and a post-mitotic quiescence program (P8) that contains self-limiting Wnt damper Nkd1 and cell cycle arrest genes Btg1/2 [34, 35].

As a generative probabilistic graphical model, the regulatory network and signal responses inferred by D-SPIN accurately reproduced both the neuronal–glial population balance and within-lineage state shifts across signaling conditions. Principal component analysis (PCA) projection of the cell population displays two major cell state clusters for glial and neuronal cell states (Figure 3C). The cell populations generated by the D-SPIN model highly resemble the experimental data distribution, including both the proportion between neuronal/glial states and the shift of density center within each cell type on the PCA projection (Figure 3D, Figure S3C). Quantitatively, the log ratio between neuronal and glial cells in the D-SPIN-generated cell population closely matches the observed log ratio across all profiled signaling conditions, with *R*^2^ = 0.993 (Figure 3E). Therefore, the inferred program-level responses by D-SPIN faithfully characterize the signal effect on the heterogeneous cell population with different glial and neuronal cell subtypes, allowing focused downstream analyses on the program-level additivity of signal actions.

The linear additivity between signal actions still holds at the gene program level. The program responses of signal combinations also lie in a three-dimensional plane as a function of two input signal levels with a scalar mid-dosage factor shared across all 14 programs (Figure 3F, Supplemental information). EGF/FGF2 and BMP4 exert opposite effects at the two displayed gene program examples, P1 and P9, and their combination effects are well-predicted by the linear sum of individual signal effects. We quantified the difference between D-SPIN-generated cell state distributions and experimental data by optimal transport (OT) distance (Figure 3G). Compared with the uniform distribution with an average distance of 8.47, the D-SPIN model with free response parameters has an average distance of 1.62, corresponding to less than 2 bit flip errors for the predicted population on average. The average distance for the D-SPIN model with additive signal responses is 1.88, only slightly higher than the model with free parameters. The results support that the additivity between the profiled signals also occurs on the gene program level, allowing the decomposition of complicated combinatorial signal effects into each signal alone.

The regulatory network model depicts a physical picture of the cell state and cell population modulation by the signal combinations. Interactions between two groups of mutually opposing gene programs collectively define a landscape for probabilities of cell states with two primary attractor basins for glial and neuronal states (Figure 3H). Each signal contributes an activating or inhibitory bias on these programs, tuning the height and location of the two attractor basins on top of the basic landscape and resulting in the abundance change between major cell types and cell subtype shifts. In signal combinations, the signal effects additively superimpose on the landscape, causing the observed population-level log additivity. Specifically for linear projections like PCA, the modulating effect on program activities can also be projected into the subspace together with the probability densities, highlighting the contribution of each gene program to the observed cell state changes (Figure 3H).

### Gene-level network model nominates regulators of neuronal-glial fate decisions

The program-level model suggested an antagonistic neuronal-glial architecture. To identify the corresponding candidate regulators that connect with signaling inputs, we constructed a directed gene-level D-SPIN network from 453 genes of transcription factors, kinases, and phosphatases expressed in the profiled population (Supplemental information) [36, 37, 38]. From the full network, we identified a 74-node core subnetwork, highlighting the signaling hubs in the fate choices between glial and neuronal cells (Figure 4A). The subnetwork is further partitioned into 7 modules, most of which correspond to a cell type component of the differentiation outcomes, including GABAergic and glutamatergic neuroblasts, glial progenitors, astrocytes, and oligodendrocytes. Among these regulators, Olig1 and Neurod6 emerge as the prominent signaling hubs with the highest number of inferred interactions.

**Figure 4.**
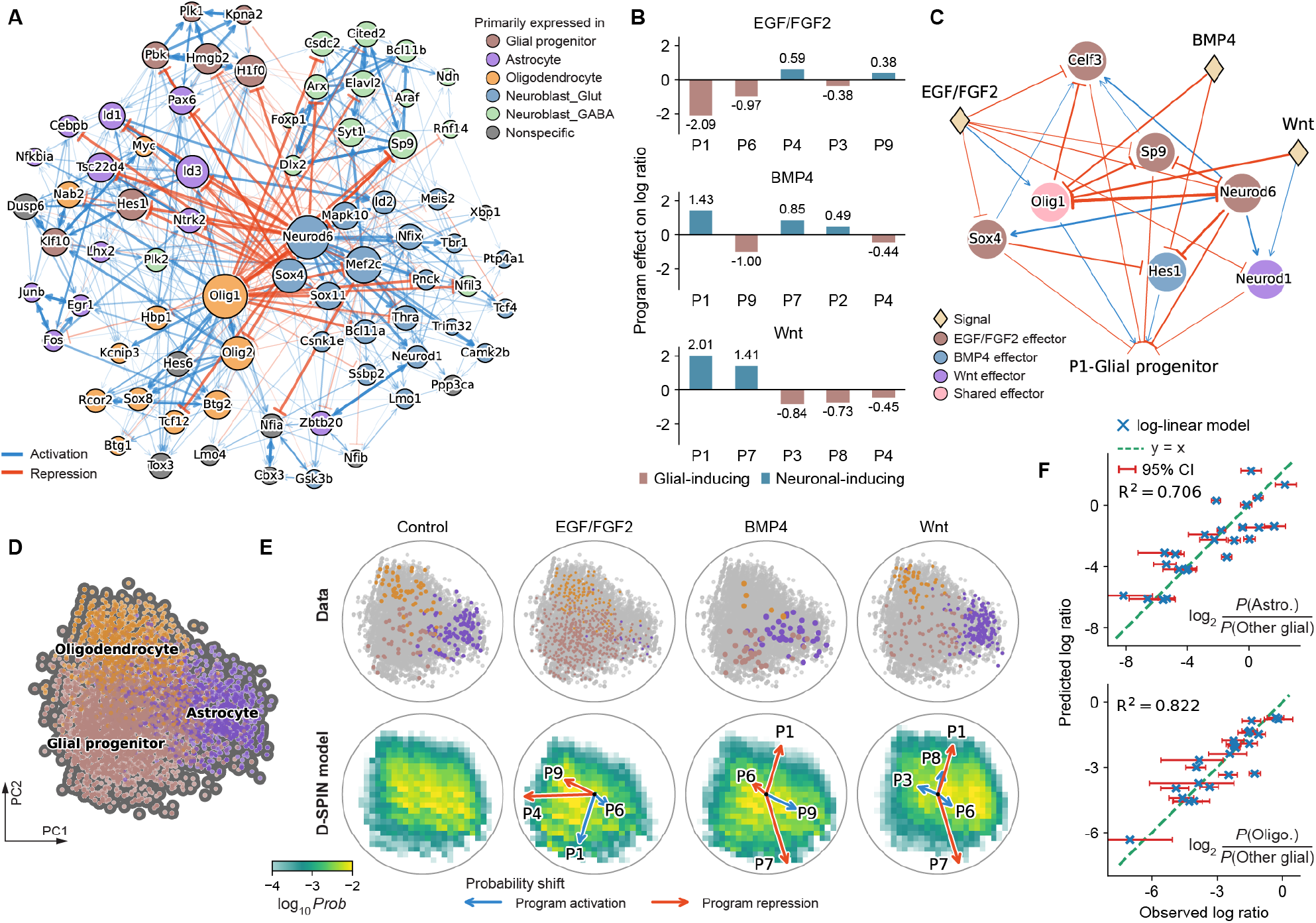
Gene-level network model nominates regulators of signature signal response programs. (A) Diagram showing the core directed subnetwork of the gene-level network model inferred by D-SPIN. Node sizes scale with the number of inferred interactions. Nodes are colored by the cell type in which the genes are primarily expressed. (B) Bar plots quantifying the contribution of program responses to the cell state shift between glial and neuronal states from counterfactual inference on the program-level network model. (C) Diagram showing the subnetwork of inferred regulators of the P1 program and the action of different signal inputs. D-SPIN nominates regulators that mediate the effect of each signal. (D) PCA projections of the program-level expression for the glial cell states only. (E) (top) PCA projections of experimental data for the glial cell population in control and signal-treated conditions. (bottom) Heatmaps of the model-generated probability landscape on the PCA projection for glial cell states. Program-level responses are also projected onto the landscape as arrows to indicate their modulation effect on probability densities. (F) Scatter plots of observed log ratios versus linear model predictions for (top) astrocytes and (bottom) oligodendrocytes compared to other glial cell states. Confidence intervals (CIs) are computed from the multinomial sampling variation of each condition.

To connect the gene regulators with the phenomenon of neuronal/glial fate switch, we performed counterfactual reasoning on the regulatory network model to identify the contribution of gene programs to cell type proportion change. We simulated cell state distributions from the model after hypothetical removal of signal effects on each gene program, and quantified the importance of programs as the neuronal/glial log ratio change after the removal (Figure S4A, Supplemental information). For all three signals, the glial progenitor program (P1) emerges as the strongest contributor to the proportion change, which contains typical early glial markers such as Fabp7, Vim, and Lgals1 (Figure 4B). Other important contributors include glycolysis (P6), oligodendrocyte program (P7), and astrocyte maturation program (P4).

We further applied D-SPIN to nominate gene regulators of P1, integrating program responses with the gene-level network to extract a core signal-to-P1 circuit (Figure 4C). The identified activating regulators are Olig1 and Hes1. Olig1 is a canonical TF in glial lineage development, and Hes1 is a highly context-dependent regulator that generally represses neuronal differentiation [39]. Inhibitory regulators of P1 includes Neurod6, Neurod1, Sp9, Sox4, and Celf3, most of which are known regulators of neuronal fate specification [40, 41]. EGF/FGF2, BMP4, and Wnt converge on this program through overlapping but distinct inferred response routes. Olig1 was inferred to respond to all three signals, providing a shared point of control. EGF/FGF2 impacts most of the targets, while Wnt also acts more selectively through Neurod1, consistent with past work on adult neurogenesis [42]. These results nominate an antagonistic switch centering Olig1 and Neurod6 as a candidate molecular implementation and suggest that additive responses between signals are achieved through a common fate-control architecture.

### Glial subtype specification follows similar log-linear models

Beyond the neuronal-glial balance, the screen also revealed signal-dependent shifts within the glial population. We then investigated whether the additive signal-integration rule extends to finer cell states. We isolated the glial states and projected the cell state distribution to the PCA directions (Figure 4D). The projection resolves three subpopulations of glial progenitors, astrocytes, and oligodendrocytes. EGF/FGF2 induces an increased glial progenitor population, while BMP4 and Wnt induce an increased astrocyte population (Figure 4E). Similarly, the program-level D-SPIN model is capable of reconstructing the cell state probability landscape that characterizes the density shift of the population. PCA projections of signal-responding programs illustrate major programs that lead to the subpopulation change, especially the cell-type-associated programs for glial progenitors (P1), astrocyte maturation (P4), and oligodendrocytes (P7).

The log-linear additive model also captures changes in glial subtype specification. Fitting the subtype outcome log ratio log_2_ *P* (Astrocyte)*/P* (Other glial) with signal input yields a reasonable fit with *R*^2^ = 0.706, while subtype outcomes of oligodendrocytes follow similar log-linear models with an *R*^2^ = 0.822 (Figure 4F). The additive rule is less accurate than the broader neuronal-glial partition. The weaker fits reflect greater sampling uncertainty from smaller subtype cell numbers, and may also indicate that finer cell type specification involves more non-additive, higher-order interaction effects between signals. The identified coefficients show that BMP4 and Wnt increase astrocyte proportion while decreasing oligodendrocyte proportion, agreeing with existing literature (Figure S4B) [43, 44]. EGF/FGF2 reduces the proportion of both fates, therefore increasing glial progenitor proportions.

Similarly, counterfactual reasoning of the network model identifies significant contributing programs to astrocyte specification (Figure S4C). Top hits are primarily cell type signature programs such as glial progenitor (P1), astrocyte maturation (P4), and oligodendrocytes (P7). Moreover, the progenitor maintenance program (P9) specifically induced by BMP4 shows the highest contribution to astrocyte fate bias. Gene-level circuits for EGF/FGF2 and BMP4 further nominate Id-family factors as candidate regulators of P9 (Figure S4D). The post-mitotic quiescence program (P8) induced by Wnt negatively contributes to astrocyte differentiation. The gene-level circuit model pinpoints three major regulators, Sp5, Lmo2, and Rcor2 (Figure S4E), with Sp5 functioning as a Wnt-induced feedback regulator and Lmo2 reported to attenuate Wnt signaling [45, 46]. These analyses suggest that distinct signals engage different regulatory modules while producing approximately additive changes in glial subtype composition.

### A annealed noise-driven switch reconciles tunable fate proportions with stable commitment

The inferred program-level and gene-level network models show signatures of opposing fate choices driven by mutual inhibitions: the program network highlights inhibitory interactions between glial progenitor (P1) and neuron differentiation (P10) programs (Figure 3A); the circuit of glial fate specification underscores the mutual inhibition between core fate regulators Olig1 and Neurod6 (Figure 4C). These mutual inhibitions are consistent with the classical model of a bistable toggle switch [47]. However, a deterministic genetic switch model explains robust commitment into alternative stable states, but does not specify how signal inputs continuously redistribute population structure. In our experiments, signal combinations do not induce an all-or-none state transition, but shift the ratio between glial and neuronal cells in a log-linear manner (Figures 2G and 2H). Nonetheless, such flexibility exhibits a constrained time window. The population cultured for 5 days with EGF/FGF2 is primarily glial cells and has lost neurogenic potential when placed into media that typically support neuronal differentiation (Figures S1G and S1H).

To reconcile these observations, we analyze a noise-driven switch model where two fate control genes *X* and *Y* mutually repress each other while each gene receives external inputs (*A, B*) and is subject to intrinsic noise (Figure 5A, Supplemental information). Without signal input, the phase portrait of the dynamical system is symmetrically divided by the separatrix into two basins of attraction for state *X* and state *Y* . A signal input that increases the production of *X* shifts the separatrix, enlarging the state *X* basin and biasing cell states towards state *X* (Figure 5B).

**Figure 5.**
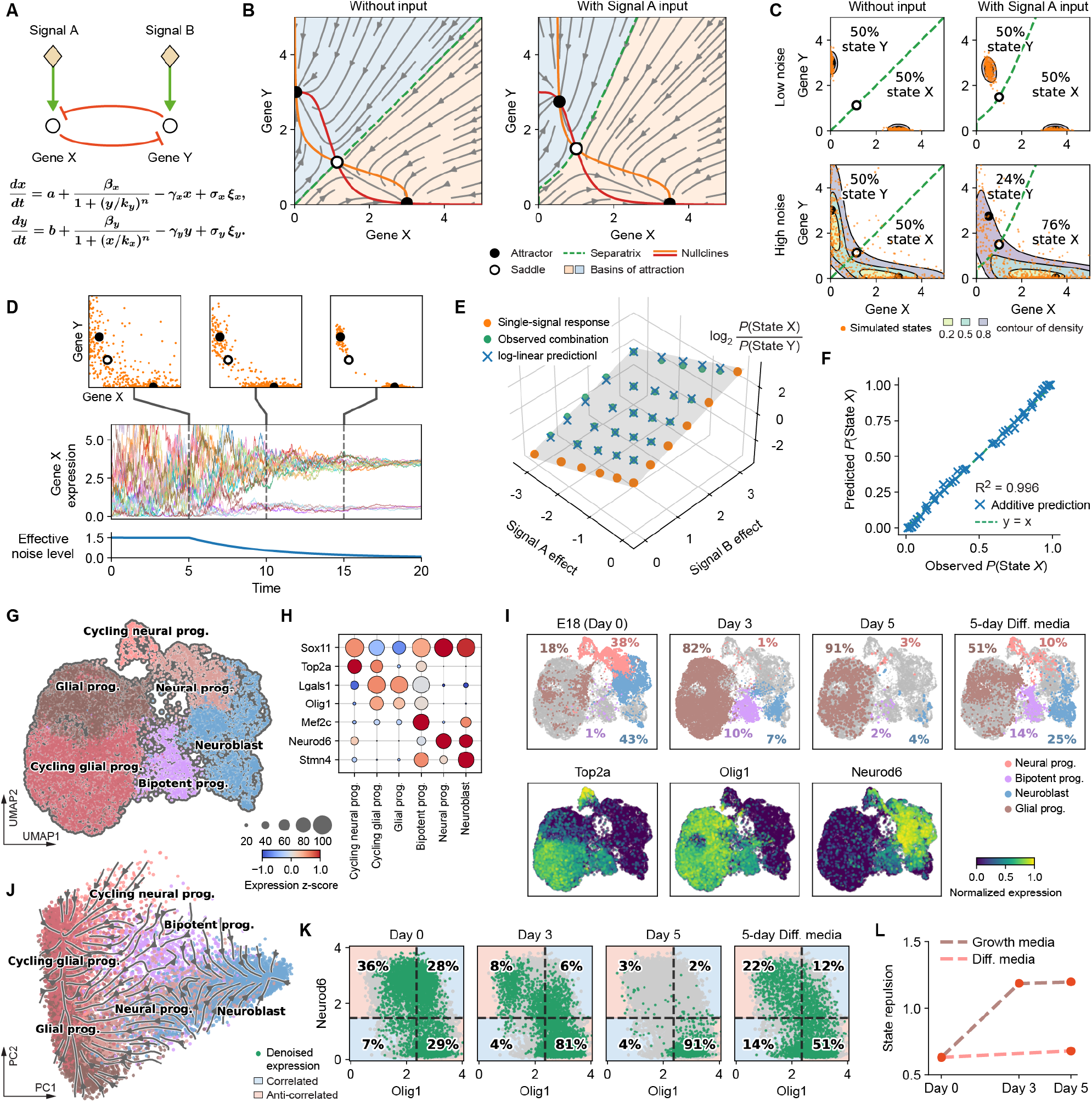
Annealed noise-driven switch model recapitulates log-linear additivity and predicts an early bipotent progenitor state. (A) Schematics and equations showing a noisy switch with external signal input. Signal inputs bias the proportion of two states, X and Y, marked by the expressions of gene X and gene Y. (B) Phase portraits of the differentiation switch with no input or signal A activation input. The phase portraits are divided into two basins of attraction of states X and Y. Signal A activation expands the basin of attraction of state X. (C) Simulated states under high/low effective noise levels and with/without the presence of signal A input, starting from an equal mixture of points near the two attractors. The signal input modulates the proportion of the two states only at a high noise level. (D) Simulated trajectories of an annealed noise-driven switch. Expression states gradually converge to the attractors with decreasing noise levels. (E) The logarithm ratios of the proportions of the two states respond to signal combinations additively with regard to single-signal responses. (F) Scatter plot showing simulated signal combination responses are well-predicted by responses of single signals. (G) UMAP embedding of a subset of single cells from early progenitor cell differentiation, including day 0, day 3, day 5 in growth media, and day 5 in differentiation media. (H) Bubble plots showing marker gene expression z-score per cell subtype. The dot sizes and colors encode the fractions of cells with detected expression and average expression levels. (I) UMAP embeddings of (top) cell populations at different culture time points and media and (bottom) expression profiles of key marker genes. The bipotent progenitor population expresses both glial and neuronal markers Olig1 and Neurod6, as well as the cell cycle marker Top2a. (J) PCA projections of the cell population with streamlines characterizing cell state transitions inferred from optimal transport. (K) Scatter plots showing denoised joint expressions of glial marker Olig1 and neuronal marker Neurod6 at different culture times and media. Cell population cultured in growth media exhibits the loss of double-positive cell states. (L) State repulsion between Olig1 and Neurod6 gene expression quantified by 0.25 log_2_ *P*_+−_*P*_−+_*/P*_++_*P*_−−_, where +, − signify the expression of the two markers.

Importantly, the effective noise level gates whether the landscape signal input induces the proportion shift between the two attractor states. We use effective noise to denote the propensity for transitions between the two attractor basins, which depends jointly on the physical noise level and the height of the transition barriers. With low noise, trajectories rarely cross the barrier between basins within simulation time, even though one attractor state is more favorable. The final state proportion remains unchanged from the initial equal mixture of attractor states (Figure 5C). On the contrary, high noise levels allow frequent barrier crossing, and signal input will continuously modulate attractor basin occupancy and induce probabilistic state switching behaviors.

Therefore, we propose an annealed noisy-driven switch model where the noise amplitude *σ* decreases over time (Figure 5D). Early high-noise dynamics permit probabilistic fate choices controlled by signal inputs, while as the noise declines, states are committed and become insensitive to signal inputs, as we observed in the differentiation experiment. Moreover, the noise-driven switch naturally reproduces the observed log-linear additivity between the two signal inputs. The log ratios log_2_ *P* (State *X*)*/P* (State *Y* ) of combinatorial inputs can be predicted by the sum of single-input log ratios, yielding accurate predictions of state proportions across a broad range(*R*^2^ = 0.996) (Figures 5E and 5F, Figures S5A and S5B).

Mechanistically, the effective noise level is not only determined by the level of transcriptional noise *σ*, but also controlled by other parameters that are more amenable to biological modulation, such as the half-saturation constant *k* and degradation rate *γ*. These factors control the barrier heights of state transitions, thus affecting the transition rate between the two attractors [48]. The noise-driven transition rate *r* between states can be characterized by the Freidlin-Wentzell quasipotential as *r* ≍ exp − Δ*S/σ*^2^, Δ*S* being the barrier height for the state transition [49]. To obtain the asymptotic behavior of the energy barrier, we analyze the model near the critical bifurcation point *β*_*c*_ of the production rate. The dynamical system has only a single stable fixed point at a low symmetric production rate *β*_*x*_ = *β*_*y*_ = *β*, which undergoes a pitchfork bifurcation to create the two stable fixed points and one saddle. Near the bifurcation, using the Landau double-well formulation and dimensional analysis [50], we obtain the asymptotic behavior *r* ≍ exp − *C*_*n*_(*β* − *α*_*c*_*kγ*)^2^*/σ*^2^*γ*, where *α*_*c*_ is the critical bifurcation point for the dimensionless equations (Supplemental information). The asymptotic behavior is further validated with numerical simulations (Figures S5C and S5D). Therefore, the results show that decreasing the effective noise level can be achieved through reducing *σ* (lower transcriptional noise), reducing *k* (stronger repression between two genes), increasing *β* (higher level of production), or decreasing *γ* (slower degradation rate), providing candidates for the search for detailed molecular mechanisms.

### Noise annealing predicts transient bipotent progenitor states and progressive fate repulsions

The annealed noisy-driven switch model indicates that probabilistic fate choices and log-additivity between signals are results of a high effective noise level in early differentiation. The model further predicts that, at early times, frequent transitions between alternative fates will create a progenitor state that exhibits signatures of both neuronal and glial cell states. To test the prediction, we isolated neuronal and glial lineages from early time points (day 0, day 3, day 5) together with cells from 5-day differentiation media culture for fine-grained cell type clustering and annotation (Figures 5G and 5H, Supplemental information).

Remarkably, we indeed identify a cell state that co-expresses neuronal marker Neurod6 and the glial marker Olig1. The cell state also expresses cell-cycle-associated genes such as Top2a, consistent with an actively proliferating progenitor state (Figure 5I). The bipotent population expresses unique markers such as Mef2c and comprises a higher proportion than neuronal states in the day 3 sample, verifying that the population does not arise from doublet artifacts (Figures 5H and 5I). The bipotent population is primarily present in the day 3 sample and the differentiation media sample with a modest proportion (10% and 14%), so it was merged with other progenitor states in earlier global analysis with more diverse cell types.

Trajectory inference and state repulsion quantification of the differentiation process are well-aligned with the annealed noise-driven switch model. We construct the cell state transition vector fields with an optimal-transport-based framework, Waddington-OT, on the denoised and batch-corrected gene expression matrix by scVI [51, 52] (Supplemental information). As visualized in the PCA projections, the vector fields contain two attractor states corresponding to glial and neuronal fates, and diverging flows at the bipotent progenitor population towards both lineages (Figure 5J). Consistent with the increasing mutual exclusivity over time in the growth media, the fraction of Olig1/Neurod6 double-positive cells declines from 28% (day 0) to 6% (day 3) and 2% (day 5) (Figure 5K). We quantify the mutual exclusivity of the two marker genes with a state repulsion index RI = 0.25 log_2_ *P*_+−_ *P*_− +_*/P*_++_*P*_−−_, which rises substantially in growth media from 0.63 to 1.20 by day 5 (Supplemental information) (Figure 5L). In contrast, the repulsion index only rises slightly in differentiation media to 0.68, indicating that the differentiation media preserve a considerable proportion of double-positive progenitor cells and therefore support a diverse progenitor population.

### Lineage-regulator mutual exclusion strengthens during *in vivo* cortical development

To assess whether the progressive segregation of alternative fate regulators observed in cell culture also applies to *in vivo* development, we analyzed a public single-cell RNA-seq dataset of mouse visual cortex development [21]. We took the subset from embryonic day 11.5 (E11.5) to postnatal day 0 (P0), filtered out non-neuronal/glial lineages and migratory lineages, and visualized the cell population with scVI batch integration and UMAP embeddings (Figures 6A and 6B, Figures S6A-S6C, Supplemental information). The dataset resolves the developmental progression from neuroepithelial cells (NEC) and radial glia (RG) through intermediate progenitors (IP) and immature neurons (IMN) to differentiated neuronal and glial lineages. We focused on two well-resolved developmental bifurcations: the separation of neuronal and glial lineages and the specification of intratelencephalic (IT) versus non-IT glutamatergic neurons.

**Figure 6.**
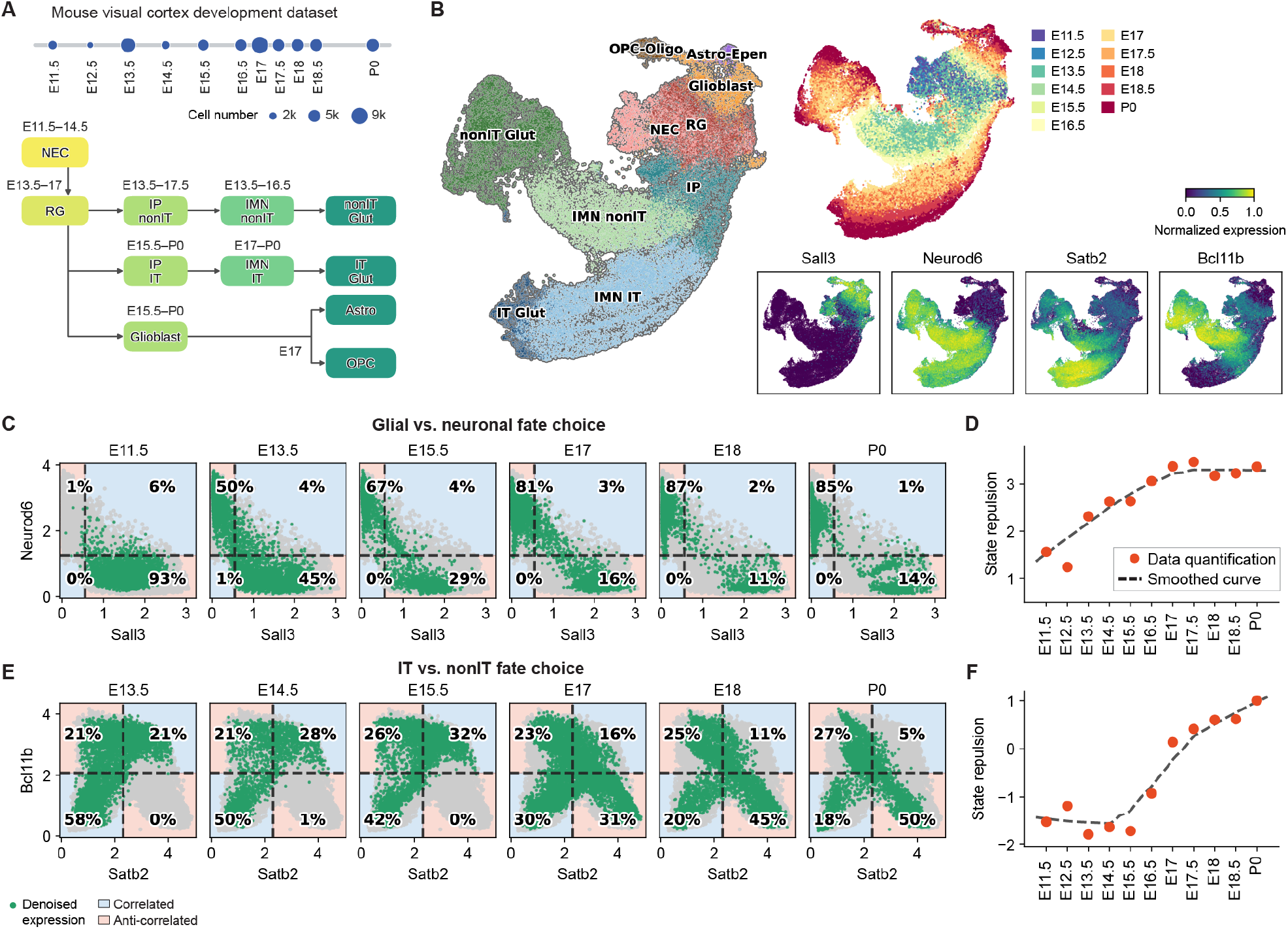
State repulsion between alternative fate regulators strengthens during mouse visual cortex development. (A) Overview of the mouse visual cortex single-cell RNA-seq dataset subset spanning E11.5 to P0 [21]. Circle size indicates the number of cells selected at each collection age. The schematic summarizes the developmental relationships between the annotated progenitor, neuronal, and glial states. (B) UMAP embedding colored by (left) annotated cell state and (upper right) collection age, together with (lower right) normalized expression of Sall3, Neurod6, Satb2, and Bcl11b as regulators for glial, neuronal, intratelencephalic (IT), and non-IT states, respectively. (C) Scatter plots showing denoised joint expression of glial regulator Sall3 and neuronal regulator Neurod6 at different developmental times. The cell population exhibits gradual loss of double-positive cell states with time. (D) State repulsion between Sall3 and Neurod6 gene expression quantified similarly to Figure 5L at individual ages. The dashed curve shows the smoothed developmental trend. (E) Scatter plots showing denoised joint expression of IT regulator Satb2 and non-IT regulator Bcl11b. (F) State repulsion between Satb2 and Bcl11b gene expression.

Gene-level network inference with D-SPIN reveals regulatory hubs for each differentiation lineage (Figure S6D, Supplemental information). We used highly-connected, lineage-associated network hubs to represent opposing branches for the two developmental bifurcations. As in the *in vitro* signal screen experiments, Neurod6 appears as the network hub for the neuronal branch. We used Sall3 to mark the glial branch, as Olig1 turns on rather late and has little expression before E17, and Sall3 is broadly expressed by the glial lineages (Figure 6B). We selected Satb2 and Bcl11b for IT/non-IT fate branches, which are established regulators for corticocortical/IT and subcerebral non-IT projection identities with known inhibitory interactions [53].

For each regulator pair, both denoised joint expression and the state repulsion index are consistent with the annealed noise-driven switch model. The fraction of Sall3/Neurod6 double-positive cells decreases from 6% at E11.5 to 1% at P0, while cells increasingly occupy either the Sall3-only or Neurod6-only state (Figure 6C). Correspondingly, state repulsion increases continuously from E11.5 through late embryonic development before reaching a plateau around E18 (Figure 6D). Similarly, the IT versus non-IT neuronal bifurcation exhibits a clear transition from regulator coexpression to mutual exclusion. Satb2/Bcl11b double-positive cells first increase from 21% to 32% as neuronal differentiation starts, then subsequently decline to 16% at E17, 11% at E18, and 5% at P0 (Figure 6E). Consistent with this transition, their state repulsion is negative during early differentiation, crosses into positive values during late embryonic development, and continues to increase toward P0 (Figure 6F).

These results provide independent *in vivo* support for the annealed noise-driven switch model. Although the two bifurcations differ in developmental timing and repulsion strength, both show progressive strengthening of mutual exclusion between alternative fate regulators. The recurrence of this pattern at different cell type hierarchies suggests that gradual fate-program segregation may be a general mechanism of population-level regulation. The early plastic states permit continuous cell type proportion modulation by combinatorial signal inputs before cell states are stably committed.

## Discussion

How progenitor populations reconcile flexible, signal-responsive fate choices with stable cell fate commitment is a central question in development. In this work, we propose an annealed noise-driven switch that achieves both, with the two behaviors separated by the effective noise level. With single-cell profiling, we show that combinations of EGF/FGF2, BMP4, and Wnt modulate the neuron-glial proportion of NPC differentiation following a simple additive log-linear model (*R*^2^ = 0.969). The responsiveness is confined to a narrow time window, as NPCs isolated from E18 SVZ lose neurogenic competence within five days in growth media culture. Regulatory network models inferred by D-SPIN reveal mutually inhibitory neuronal and glial modules, motivating the noise-driven switch model. High effective noise permits transitions between attractor basins, so signal inputs can tilt the landscape continuously to redistribute the population. As noise declines, transitions are suppressed and the population composition is fixed. The model further predicts early progenitor states with regulators of both fates, which we verify in our experimental data and in a public dataset of *in vivo* visual cortex development.

The transcriptomic resolution of single-cell profiling combined with D-SPIN regulatory network inference provides mechanistic insights beyond the proportion change from cell type counting. Log-linear proportions could arise from the modulation of exponential growth rate, fate choice probabilities at differentiation, or chances of survival. As inferred by the D-SPIN model and also visualized in the expression heatmap (Figures 3A and 3B), the mitosis program (P5) shows little change across signaling conditions, and apoptosis was not identified as a significant activity in the program discovery pipeline, whereas signals primarily target lineage-specifying programs (P1, P4, P7). Counterfactual inference also identifies glial progenitor program P1 as the dominant contributor to proportion change under all three active signals. These results suggest that population restructuring is achieved primarily by biasing fate during differentiation rather than by altering growth or survival, agreeing with prior observations [43]. Moreover, signal regulations mostly impinge on glial-associated programs, and the balance between excitatory versus inhibitory neurons is not significantly shifted by most signals except EGF/FGF2, suggesting that signals mainly modulate the probability of adopting glial fates during progenitor differentiation (Figures S4F-S4H).

During development, progenitors execute coordinated fate choices under combinatorial signaling environments and morphogenic gradients, and the mechanisms of decoding this information remain an open question. Our findings point to a simple additive signaling code. When the transitions between alternative states are frequent, the ratio between different fates is set by the difference in barrier heights separating them. Signal inputs shift heights of individual barriers, making additive combination in log-ratio space the leading-order behavior of the system. We observed stronger additivity for the broader neuronal/glial lineages (*R*^2^ = 0.969) than for the finer specification of astrocytes or oligodendrocytes (*R*^2^ = 0.706, 0.822). The scale dependence suggests that coarse-graining may reinforce the additivity, as nonlinear cross-talks are buffered when aggregated into expression programs and major cell types, even if underlying genes interact in a more complex, context-dependent manner.

Prior work has shown that combinations of signal ligands can have complex, non-additive properties, and certain developmental outcomes require precise combinations between signals, such as the generation of mesoderm cells and somitogenesis [54, 14, 55]. Nonetheless, our results highlight a complementary regime where signals combine additively: both cell proportion log ratio and program-level responses are well captured by a linear model. Additivity in biological signal processing is considered a general principle that appears to be the default mode of computation [54, 56, 20]. Systematically extending this code across a broader collection of signals and other progenitor populations could enable selective expansion or depletion of target cell states within heterogeneous populations, providing a strategy for directing stem cell fates for therapeutic applications.

Progenitor states co-expressing regulators of alternative lineages have been described across systems as multilineage priming [57, 58, 10]. Although whether such co-expression is required for cell fate decision remains debated [59], our results provide a candidate functional role of these states. Frequent transitions between attractor states create a responsive window to signal inputs, allowing continuous tuning of lineage proportions. We identify a transient Olig1/Neurod6 double-positive progenitor state, with the two markers nominated by the D-SPIN network as opposing regulatory hubs. In the developing visual cortex, state repulsions between both Sall3/Neurod6 and Satb2/Bcl11b progressively increase from E11.5 to P0. These observations extend multilineage priming to a dynamic perspective, where increasing state repulsion indicates the transition from plasticity to commitment.

In the noise-driven switch, the effective noise refers to the propensity for transitions between attractor basins, and depends on both the amplitude of expression fluctuations and the height of the barrier. At the molecular level, transcriptional noise can be amplified by altering DNA topology to temporarily impede transcription [60]. Heterogeneity can also arise dynamically rather than stochastically: delayed self-inhibition, as in Hes1, generates sustained oscillations that diversify gene expression states within a uniform environment [9]. Through theoretical derivations, the barrier height can be modulated by regulatory binding affinity, production rate, and degradation rate. The binding affinity can be modulated by forming complexes with other transcription factors and cofactors. Both decreasing the production rate and increasing the degradation rate cause reduced gene expression levels, while advances in metabolically labeled single-cell RNA profiling enable capturing nascent transcript dynamics and distinguishing these scenarios [61].

The Waddington landscape metaphor indicates that cell states are robustly constrained within valleys upon external stimulation, while our findings suggest a quantitative refinement. These valleys are also tunable by signal input to allow cell types to continuously traverse different states. Beyond changing cell type abundances, signals shift the state of each cell type, preserving core lineage programs while turning on the expression of signal-specific response programs [62]. Despite the nonequilibrium nature of these systems, D-SPIN, as a statistical physics model, captures the continuous distribution shift of program-level states with low quantitative error, demonstrating the utility of such abstractions to model complex biological processes [63].

## Data availability

Supplementary tables of the analyses are available at Caltech Research Data Repository https://doi.org/10.22002/dtmcr-qc483

Gene counts, metadata, and preprocessed single-cell data of all profiling experiments, as well as inferred D-SPIN models, are available at Caltech Research Data Repository https://doi.org/10.22002/471a4-2vz62

## Author Contributions

Conceptualization, J.J., S.C., M.T.; Methodology, J.J., S.C., M.T.; Data Analysis, J.J., S.C., V.Y., P.R., M.T.; Experimental Investigation, S.C., J.H.P., T.T.; Resources, S.C., M.T.; Writing, J.J., S.C., M.T.; Visualization, J.J., S.C., M.T.; Supervision, J.J., S.C., M.T.; Funding Acquisition, M.T.

## Acknowledgments

We would like to thank Eric Siggia, Michael Elowitz, Lior Pachter, Zev Gartner, Eric Chow, Yaron Antebi, Allan-Hermann Pool, David Schaffer, Carlos Lois, Chris McGinnis, and Elliott Robinson for feedback and discussions; Jase Gehring for helping with the ClickTag experiments and discussions; Inna-Marie Strazhnik for figure illustrations; Sandy Nandagopal, Elisha Mackey for reagents and cells; members of the Thomson Lab; and the Beckman Institute Single-cell Profiling and Engineering Center (SPEC). Funding support for this project was provided by the NIH Office of the Director (5DP5OD012194-04), the Shurl and Kay Curci Foundation, and the Heritage Medical Research Institute.

## Declaration of interests

MT is a co-founder of CognitiveAI and YurtsAI.

## Supplementary Figures

**Figure S1.**
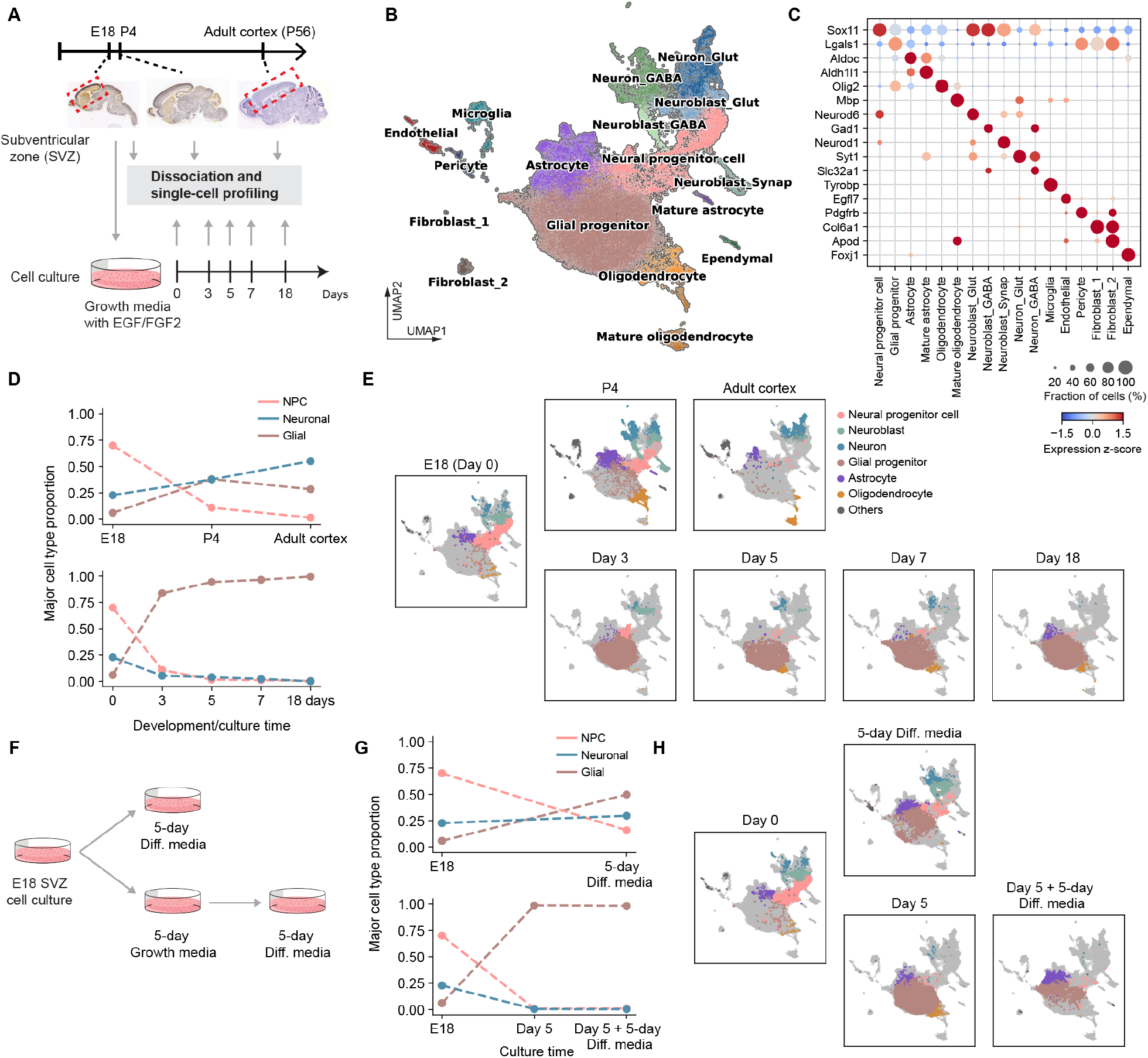
Single-cell profiling reveals gradual restriction of population diversity and neurogenic potential during cell culture; related to Figure 2. (A) Experimental schematics showing time points when cells were collected and profiled from (upper) the developing mouse brain and (lower) cell culture. Cell cultures were initiated from cells dissociated from the E18 subventricular zone (SVZ). (B) UMAP embedding of 60k quality-controlled single cells from the developmental reference and culture time-course profiling experiments. The cell population contains glial and neuronal cells from progenitor to mature states, and other brain-associated cell populations. (C) Bubble plots showing marker gene expression across annotated cell types. Dot size indicates the fraction of cells within each cell type with detected expression, and dot color indicates average expression level. (D) Line plots showing the proportion changes of neural progenitor cells (NPCs), glial and neuronal cells during (upper) developmental stages or (lower) culture timepoints in growth media. *In vivo* development generates diverse neuronal and glial populations, whereas cell culture rapidly concentrates the population in glial states. (E) UMAP embeddings of the cell population at different developmental stages or culture timepoints. (F) Schematics of experimental design to test whether prior expansion of the glial population in cell culture restricts subsequent neurogenic potential. (G) Line plots showing the cell type proportion changes after (upper) culturing directly in differentiation media and (lower) culturing in growth media, then differentiation media. Immediate differentiation generated both neuronal and glial populations, whereas cells previously expanded in growth media remained glia-dominant after transfer to differentiation media. (H) UMAP embeddings of the cell population directly cultured with differentiation media or first cultured with growth media.

**Figure S2.**
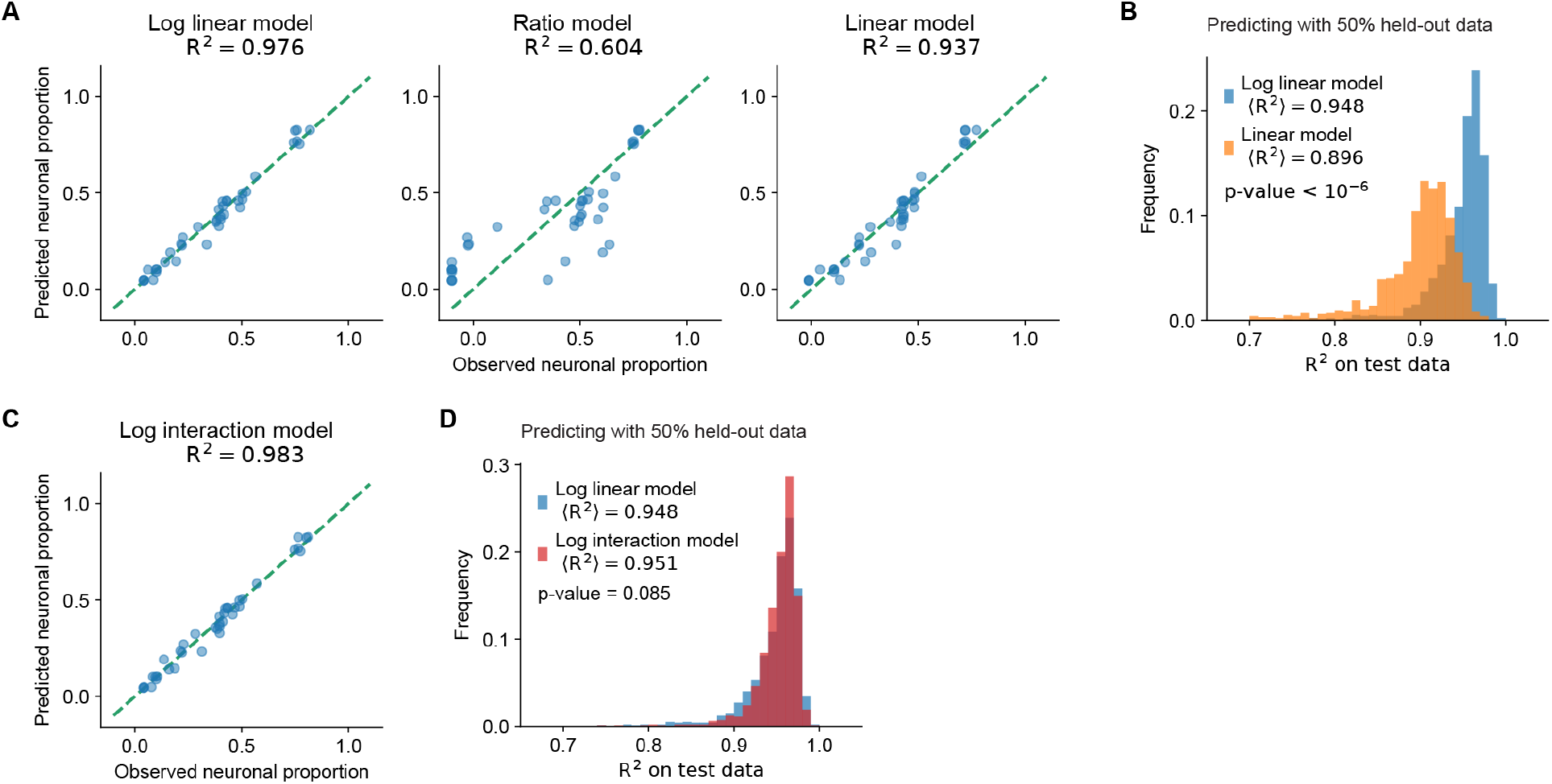
Log-linear model outperforms other alternative models; related to Figure 2. (A) Scatter plot shows predicted neuronal proportion *P* (Neuronal) with different regression models by fitting (i) log_2_ *P* (Neuronal)*/P* (Glial) (log linear model) (ii)*P* (Neuronal)*/P* (Glial) (ratio model) (iii) *P* (Neuronal) (linear model). The log-linear model achieves the best performance. (B) Histogram quantifying the *R*^2^ of predicting *P* (Neuronal) with held-out data for the linear model and the log-linear model. The model is fitted with 50% of the data and evaluated on the other 50% as test data. (C) Scatter plot shows predicted *P* (Neuronal) with pairwise signal combinations (interactions), such as BMP4 + Wnt, included as extra predictors. The *R*^2^ of fitting only shows marginal improvement. (D) Histogram quantifying the *R*^2^ of predicting *P* (Neuronal) with 50% held-out data for the log interaction model. The prediction accuracy does not significantly differ (p=0.085) from the log-linear model.

**Figure S3.**
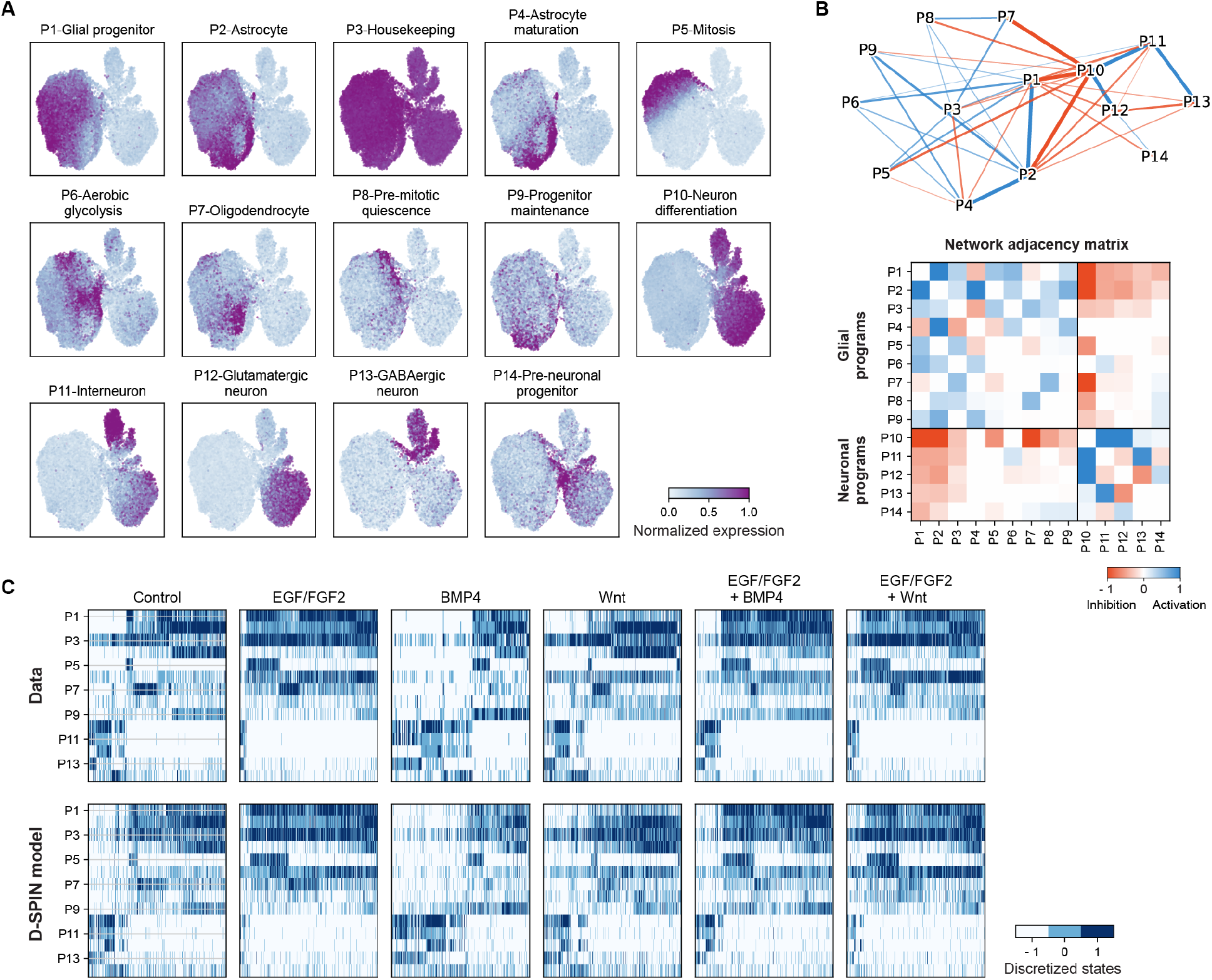
Program-level network model reconstructs cell state distributions across signaling conditions; related to Figure 3. (A) UMAP rendering of the normalized expression level of each gene program. Each gene program is specifically expressed in cell states localized in regions on the UMAP. (B) (top) Network diagram of the program-level regulatory network model inferred by D-SPIN. The network is partitioned into two modules that mostly contain glial/neuronal-associated programs. (bottom) Heatmap showing the adjacency matrix of the inferred network. The interactions are largely activating within each module and inhibitory between the two modules. (C) Heatmaps of program expression for (top) the observed data and (bottom) 10,000 random samples generated by the D-SPIN model. The D-SPIN-generated transcriptional states are well-aligned with each cell cluster structure in the data heatmaps.

**Figure S4.**
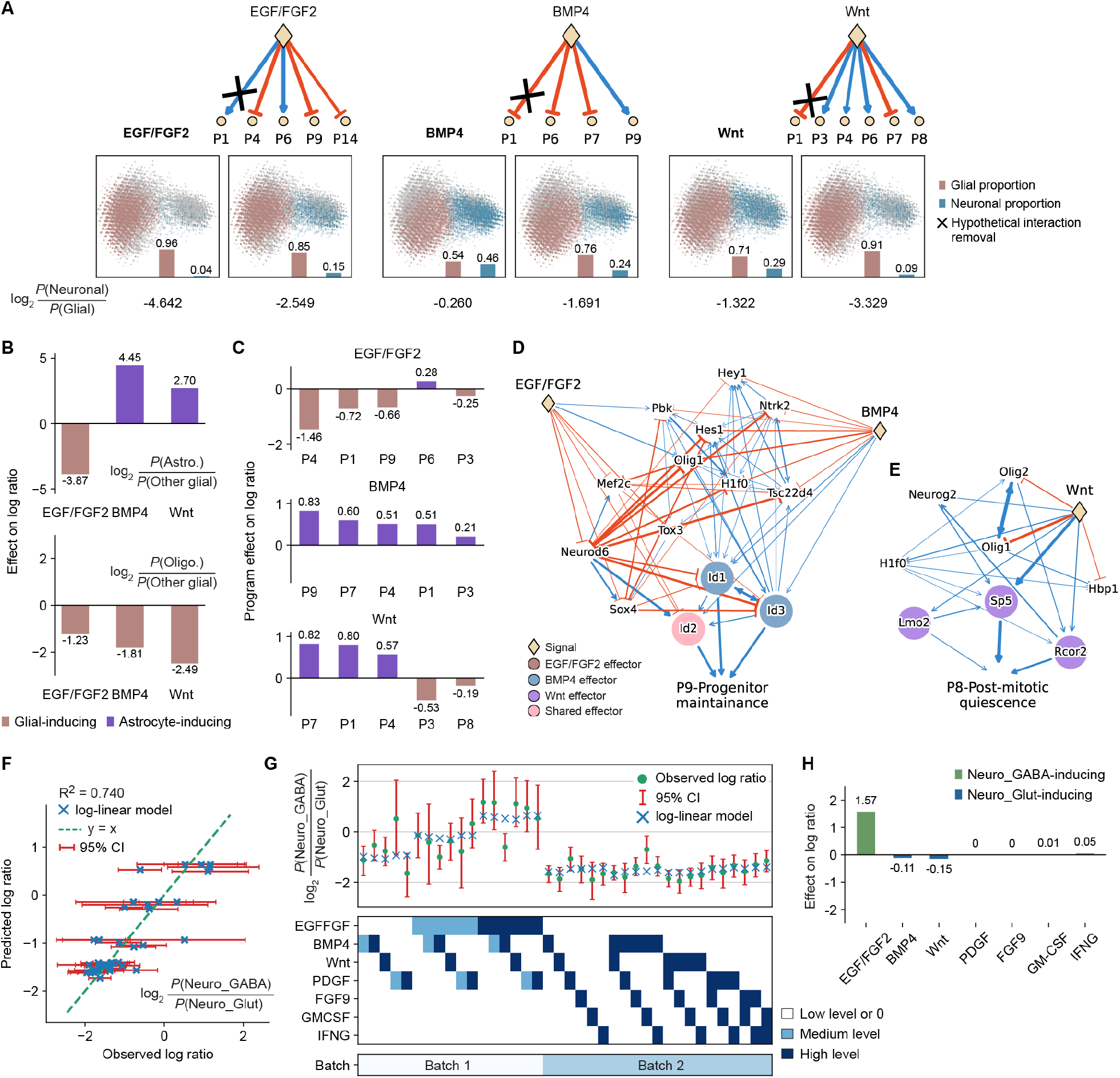
Counterfactual reasoning for neuronal/glial fate changes; log-ratio changes for glial and neuronal subtypes; related to Figure 4. (A) (top) Network diagrams and (bottom) PCA projections of D-SPIN-generated cell state distributions before and after hypothetical removal of signal effects on P1. The changes in the log ratio after removing P1 response quantify the contribution of P1 to the neuronal/glial fate choices. (B) Coefficients of the linear model on the response of glial cell type proportion log ratios for (top) astrocytes and (bottom) oligodendrocytes. (C) Bar plots quantifying the contribution of program responses to the glial cell states shift towards astrocyte states. EGF/FGF2 and BMP4 have a signature program P9, and Wnt has a signature program P8. (D) Diagram showing the subnetwork of inferred regulators of the P9 program and the action of EGF/FGF2 and BMP4 signals. (E) Diagram of P8 program regulators and Wnt signal action. (F) Scatter plot of the observed logarithm of the ratio between GABAergic neuroblasts and glutamatergic neuroblasts versus predicted log ratios by a linear model. The 95% confidence intervals (CIs) are broad for some samples as the total number of neuronal cells in these conditions is small. (G) The log ratios between neuroblast types show little change with signal inputs except for EGF/FGF2 across the profiled signaling conditions. Concentrations of each signal are normalized to up to three levels. (H) Coefficients of the linear model on the response of neuroblast type log ratios. Only EGF/FGF2 has a significant effect, increasing the proportion of GABAergic neuroblasts.

**Figure S5.**
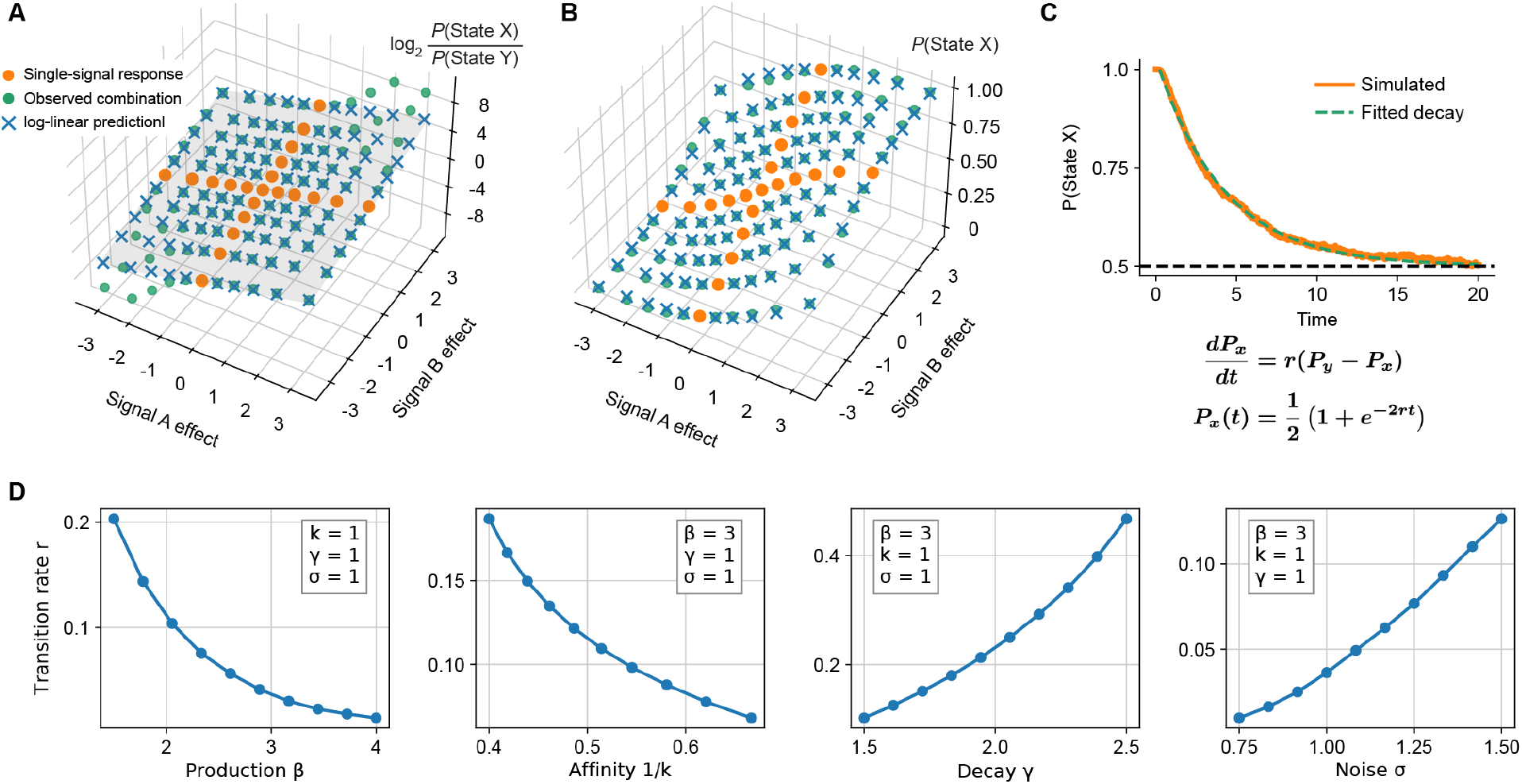
Log-additivity of annealed noisy toggle switch; transition rate between attractors; related to Figure 5. (A) The logarithm ratios of the proportions of the two states respond to signal combinations mostly additively with regard to single-signal responses. Deviations from additivity occur at log ratios with large magnitude, which have less impact on predicting the probability. Log ratios are clipped between [-10, 10] for visualization. (B) Log-additive model on single-signal responses predicts signal combination effect well. The non-linear deviations in the large-magnitude log ratios do not affect predictions as the probability is close to 0 or 1. (C) Examples of simulating transition rates *r* between two attractor points. The proportion of State X decays with time exponentially following a simple reaction rate model. (D) Numerical evaluations of transition rates as a function of model parameters. As predicted by the asymptotic analysis, the transition rate between attractors, or equivalently, the effective noise levels, decreases with production rate and binding affinity and increases with decay rate and physical noise level.

**Figure S6.**
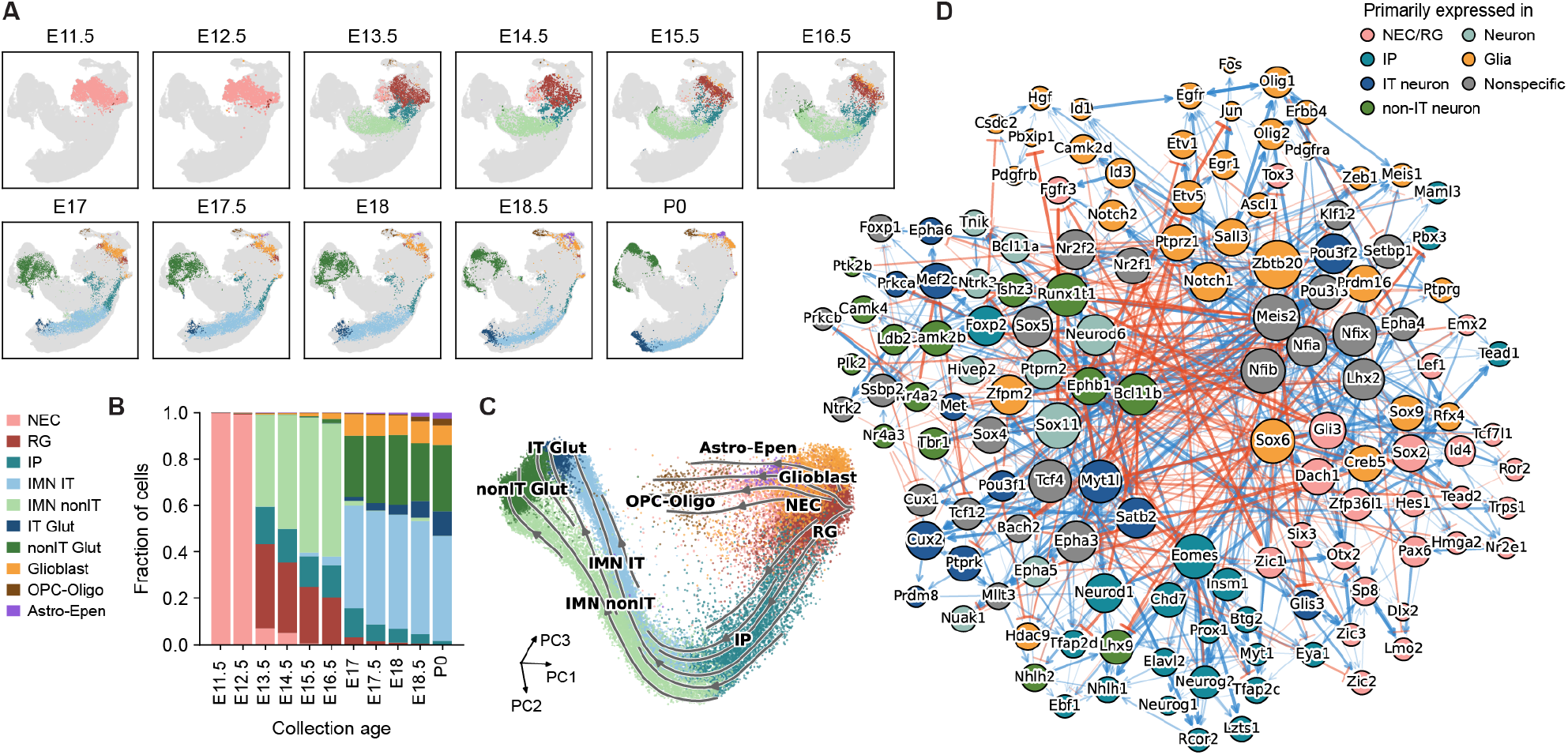
Cell-state organization and regulatory network inference of the mouse visual cortex development dataset; related to Figure 6. (A) UMAP embeddings time course of the mouse visual cortex developmental dataset from embryonic day 11.5 (E11.5) to postnatal day 0 (P0). (B) Bar plots showing the relative abundance of annotated cell states among the cells profiled at each collection age. (C) PCA projections of the cell population with streamlines characterizing cell state transitions inferred from optimal transport. (D) Diagram showing the core directed subnetwork of the gene-level network model inferred by D-SPIN from the cortex development dataset. Node sizes scale with the number of inferred interactions. Nodes are colored by the cell type in which the genes are primarily expressed.

## Method details

### Experimental Model and Subject Details

The animal study involving the dissection of the adult mouse brain cortex was reviewed and approved by the Caltech Institutional Animal Care and Use Committee (IACUC). All other developmental time points involved mouse embryos (C57BL/6 purchased from a commercial supplier, Brainbits Inc) whose animal protocol was approved by the Southern Illinois University School of Medicine Laboratory Animal Care and Use Committee.

### Mouse brain tissue dissociation

Tissue from mouse brain (C57BL/6) was purchased from Brainbits or acquired from collaborators (Allan-Hermann Pool, Oka Lab, Caltech; Elisha Mackey, Gradinaru Lab, Caltech), and digested using 6 mg of Papain (Brainbits) at 2 mg/mL in Hibernate E medium (Brainbits) for 30 minutes at 30°C. After incubation, tissue was triturated for 1 minute using a silanized Pasteur pipette. Dissociation medium containing the dissociated cells was centrifuged for 1 minute at 200x g, and the cell pellet was either resuspended in media for further culture or resuspended in 1x PBS + 0.04% BSA for single-cell profiling through the 10X Genomics platform.

### Subventricular zone cell derivation

Neural progenitors are dissociated from the combined cortex, hippocampus, and ventricular zone of E18 mice (Brainbits). After dissociation and resuspension of the cells in NPgrow media (Brainbits), cells were counted and plated onto ultra-low attachment 100 mm culture plates (Corning) at one million cells per plate in 10 mL of NPgrow, and supplemented with 1% Penicillin-Streptomycin (Gibco). NPgrow media consists of Neurobasal media, 2% B27 (minus Vitamin A), 0.5 mM Glutamax, 20 ng/mL EGF, and 20 ng/mL FGF2. Cells were cultured at 37°C and 5% CO2, and the supplemented medium was changed every 2 to 3 days.

### *In vitro* differentiation with media

*In vitro* expanded and directly dissociated progenitor cells were differentiated for 5 days in custom Brainbits NbCo-culture media that support the growth of both neuronal and glial differentiated cells. The media consisted of Neurobasal, 2% B27 (minus Vitamin A), 0.5 mM Glutamax, 10% FBS, and 1% Penicillin-Streptomycin (Gibco), and was supplemented with a proprietary mixture of creatine, estradiol, and cholesterol. Cells were seeded onto poly-ornithine/laminin coated plates at a density of 50k cells/cm^2^, and cultured at 37°C and 5% CO2. Medium was changed every 2 to 3 days. Cells were cultured for 5 days before dissociation for single-cell profiling.

### Combinatorial signaling screen

We applied combinatorial signals to cells dissociated from the SVZ region of the E18 mouse brain to understand how signals regulate the population structure and transcriptional state of progenitor cells. We used NPGrow media, which did not contain EGF/FGF2, and added signaling factors to the final concentrations given in the SI Table. For EGF, FGF2, BMP4, PDGF-AA, and FGF9, the maximum concentration used was 20 ng/mL. For IFN-G and GM-CSF, the maximum concentration used was 3 ng/mL, and for CHIR99021, the maximum concentration used was 3 uM. After cells were resuspended in media, they were cultured for 5 days before dissociation for ClickTag multiplexing and single-cell profiling.

### Experimental multiplexing using ClickTags

Cultured cells were dissociated in Accutase into a single-cell suspension. Cells were then fixed in ice-cold 80% methanol. Prior to cell labeling, two methyltetrazine (MTZ) activated barcoding oligos were combined for each labeling sample, NHS-trans-cyclooctene (NHS-TCO) was then added and left to reaction for 5 minutes. The previously fixed cells were then mixed thoroughly with the barcode oligo solution and incubated for 30 minutes at room temperature. The labeling reaction was then quenched with the addition of Tris-HCl and methyltetrazine-DBCO. After the quenching step, cells from all labeling samples were pooled together along with the addition of two volumes of PBS-BSA, then centrifuged at 800x g for 5 minutes, followed by two more cycles of PBS-BSA washes and centrifugation steps. After the final centrifugation step, the cell pellet was resuspended in 100 uL of PBS-BSA and counted on a hemocytometer. Cells were then loaded onto 3-4 lanes, targeting 10,000 cells per lane, of the Chromium Controller and processed as recommended by the 10x Genomics v2 Single Cell 3’ reagent kit protocol until completion of the cDNA amplification PCR step. At this step, SPRI size-selection was used to separate the barcode oligos from the cDNA, which were then processed separately. We used a dual-tagging approach, so that each experimental sample gets two ClickTag barcodes.

### Barcode demultiplexing

Filtered cell barcodes from the aligned gene expression (Cellranger) are used as a whitelist to parse ClickTags data from FASTQs. Tag sequences extracted from fastqs are aligned to a list of known ClickTag sequences, allowing for 1 mismatch (Levenshtein distance ≤ 1). Counts for ClickTags are accumulated and stored in a cell barcode by tags matrix. To threshold the data, we computed a count distribution for each individual ClickTag, and used Otsu’s method to obtain optimal thresholds for each ClickTag sequence. Each cell is thresholded for each barcode independently, and cells that have the correct barcode pair combinations are retained.

### Single-cell data processing, clustering and visualization

Single-cell profiling data obtained from different experiments were grouped into three major datasets, referred to as Brain Development, Signal Response, and Early Differentiation. The Brain Development dataset contains profiling of *in vivo* tissues of E18 SVZ, mouse brain at postnatal day 4 (P4), and adult cortex, as well as *in vitro* cell culture of neural progenitor cell (NPC) population with growth media and differentiation media. The Signal Response dataset contains two batches of multiplexed single-cell profiling of a total of 40 dosage combination signal treatment conditions. The Early Differentiation is a subset of the Brain Development dataset with only day 0, day 3, and day 5 in growth media and day 5 in differentiation media. The three datasets are processed separately to highlight unique biological variations within similar experimental setups.

The three datasets undergo the same quality control with the following steps:

1. Remove cells that have fewer than 200 genes expressed.
2. Remove genes that are expressed in fewer than 10 cells.
3. Remove cells in which more than 20% of transcripts are mitochondrial genes.

Due to the variability of captured transcript number for each cell, gene expression data is normalized for each cell to a total target count of 10^4^. Due to measurement noise, genes that have constant expression across cells do not contribute to biological impact, but only increase noise in the data. Therefore, single-cell data analyses are performed on a subset of genes that have different expression levels across cells, i.e., highly variable genes. We used a customized coefficient-of-variation (CV) filtering for the negative binomial distribution as developed in our previous work [1]. For each dataset, we selected 3,000 highly variable genes according to the rank of negative binomial CV filtering and conditioned on having nonzero expression in more than 2% of the cells. Selected highly variable genes are log-normalized by the transformation log(*x* + 1). All the following analyses were performed on the log-transformed data except otherwise noted.

To cluster and visualize single cells, we corrected the batch effect by constructing latent representations with scVI on raw gene counts for each dataset separately with similar setups (dispersion=‘gene-batch’, n latent=10, n layers=2, batch size=2048, max epoch=200 (Brain Development) or 400 (Signal Response and Early Differentiation)) [2]. The training epoch number difference is due to different cell numbers in the three datasets. We constructed a 30-nearest-neighbor graph (scanpy.pp.neighbors) on the latent representation, computed Leiden clustering for cell clusters (scanpy.tl.leiden), and computed UMAP (scanpy.tl.umap) for visualizations. For the Early Differentiation dataset, after the first round of Leiden clustering, clusters that were neither glial nor neuronal lineages, such as endothelial and microglia, were removed, and we redid the neighborhood graph construction, Leiden clustering, and UMAP embedding computation.

### Regression and confidence intervals for cell proportion logarithm ratios

To quantify the fluctuations of cell proportion log ratios due to the finite cell number in each condition, we performed bootstraps to compute confidence intervals. For each condition with *N* total cells and *n*_1_, *n*_2_ two cell types of interest, we sample from the multinomial distribution

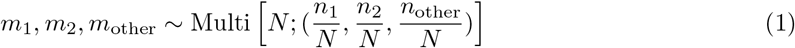

for 10^5^ times, and use the 2.5 and 97.5 percentiles of log(*m*_1_ + 0.5)*/*(*m*_2_ + 0.5) as the 95% confidence intervals. The extra 0.5 term is Laplace smoothing to avoid 0s in the logarithm. We used ℓ_1_-regularized linear regression (Lasso) on cell proportion log ratios with sklearn.linear model.Lasso (alpha = 0.005). Signals with three dosages were set to 0, 0.5, and 1 as predictors, and signals with two dosages were set to 0 and 1 as predictors.

### Gene program discovery

Gene programs were identified with the D-SPIN program discovery pipeline [1]. D-SPIN uses orthogonal nonnegative matrix factorization (oNMF) to identify coexpressing and mutually independent gene programs. oNMF is similar to principal component analysis (PCA), but with the extra constraints that gene coefficients in each component are nonnegative and that each gene can only belong to a single component. Formally, oNMF solves the following optimization problem

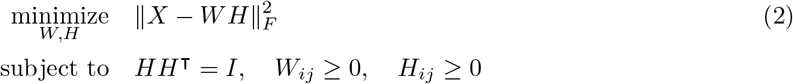

where *H* ∈ ℝ^*K×n* gene^ is the gene program representation, *W* ∈ ℝ^*n* cell*×K*^ is the cell state represented on the gene programs and 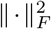 is the matrix Frobenius norm. *HH*^⊺^ = *I* ensures all program components are non-overlapping.

We used the Python package dspin to identify 25 gene programs with DSPIN.gene program discovery (num repeat=100, cluster key=‘leiden’, params= {‘onmf epoch number’: 2000} ). Specifically, as the solution of oNMF depends on random initialization, the parameters request the program to run oNMF with 100 different random initializations, each random seed to iterate for 2000 epochs, and to summarize consensus gene programs as results. The program number 25 is determined by manually examining the quality of identified gene programs from the decomposition with 15, 20, 25, and 30 programs, especially whether the signal-specific program for BMP4 and Wnt is recovered.

The identified 25 gene programs were further merged into 14 gene programs by combining programs that have similar expression patterns across cell subtypes, especially programs characterizing different neuron activities that are irrelevant to the analysis. The program merging was performed by DSPIN.gene program discovery(prior programs=prior gene list), which computes the optimal weighted average of gene expression in each program with the lowest reconstruction error.

### Regulatory network model constructions with D-SPIN

The D-SPIN framework infers a regulatory network model of cell state distributions by solving a context-dependent inverse Ising problem [1]. D-SPIN formulates the distribution of the three-way discretized transcriptional state ***s***, *s*_*i*_ ∈ {− 1, 0, 1} as the effect of a constant pairwise interaction matrix ***J*** and condition-dependent local activities ***h***^(*n*)^, (*n*) indexing for experimental conditions. Formally, the probability distribution is

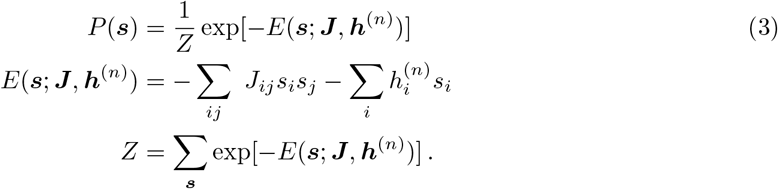

***J***, ***h***^(*n*)^ are solved from experimental observations by maximizing the log likelihood function or its surrogate

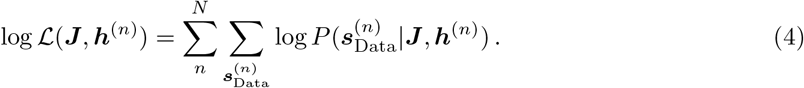

We used the Python package dspin to solve the inference problem. The program-level network was constructed with DSPIN.model program.network inference(sample id key=‘sample batch’, method =‘maximum likelihood’, params=‘stepsz’: 0.2, ‘num epoch’: 1000). The exact maximum likelihood inference was used as the distribution of 14-node networks can be explicitly computed to ensure optimal accuracy.

For gene-level network construction, we selected regulatory genes, including transcription factors (TFs), kinases, and phosphatases from databases [3, 4, 5]. We filtered for genes that are expressed in more than 2% of the cells, resulting in 314 transcription factors (TFs), 98 kinases, and 41 phosphatases. The 453-node network was constructed with DSPIN.network inference(sample id key=‘sample batch’, method=‘pseudo likelihood’, directed=True, params=‘stepsz’: 0.04, ‘num epoch’: 1000). The inference used pseudo-likelihood, which is the only method that scales to large networks with high accuracy and provides information about interaction directionality. We associated the 14 gene programs with regulators in the gene-level network using the regulatory discovery algorithm in D-SPIN DSPIN.model gene.program regulator discovery(program representation, params=‘num epoch’: 10000, ‘stepsz’: 0.05)

### Regression model of D-SPIN-inferred responses

In the regression of cell proportion log ratios, the middle dosage level was set to 0.5 in the predictors, as the dependent variable (log ratios) is a 1 one-dimensional scalar. Therefore, allowing flexible dosage response might cause overfitting. In the regression of program-level responses, the 14 gene programs provide enough information content to quantify the effect of middle dosage as the full dosage response scaled by a factor *γ*, 0 ≤ *γ* ≤ 1.

There are three signals that have three dosages, EGF/FGF2, BMP4 and PDGF; so we introduced three scaling factors *γ*_*e*_, *γ*_*b*_, *γ*_*p*_ for the middle dosage of each signal. The regression is now formulated as an optimization problem

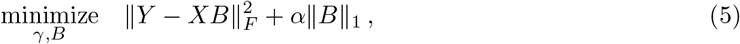

where *Y* = [***h***^(1)^, ***h***^(2)^, …, ***h***^(*N*)^]^⊺^ is the D-SPIN inferred response for all conditions and programs as the regression target; *B* ∈ ℝ^*n* signal*×n* program^ is the inferred signal-specific responses; *X* ∈ ℝ^*n* condition*×n* signal^ is the design matrix.

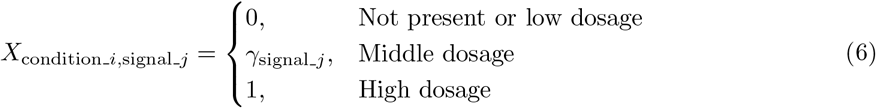

The inferred parameters are *γ*_*e*_ = 0.607, *γ*_*b*_ = 0.285, *γ*_*p*_ = 1.00. PDGF has little effect on the cell population; *γ*_*p*_ = 1 also suggests that the middle dosage of PDGF does not exhibit significant differences from the full dosage. The regression on D-SPIN gene-level responses also follows the same formulation, with the *γ* factors fixed as the program-level responses to ensure consistency.

### Simulating cell states from D-SPIN model and counterfactual inference

The inferred regulatory network and signal responses define a cell state distribution in the space of program expressions by Eq. 3. We first explicitly compute the distribution on 3^14^ ≈ 4.8 *×* 10^6^ states, then sample 10^6^ samples from the distribution with numpy.random.choice. To assign the neuronal/glial cell type labels to sampled states, we trained a linear binary classifier (support vector machine, SVM) on the experimental data with sklearn.svm.LinearSVC(C=0.01, penalty=‘l1’), and used the linear classifier to label glial states. To assign astrocyte/oligodendrocyte subtype labels, we trained SVMs for each subtype, and labeled astrocyte/oligodendrocyte by the intersection of the glial classifier and the subtype classifier.

In the counterfactual inference, given the network ***J*** and response ***h***^(*n*)^ for condition *n*, baseline activity for control sample ***h***^(0)^, the counterfactual response on program *k* is defined as

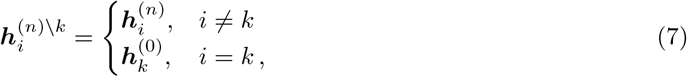

which means setting the response on program *k* to the activity level of the control sample 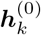. For any observable Φ defined on the distribution *P* (***s***; ***J***, ***h***^(*n*)^), for example, the log ratio between neuronal states and glial states, the counterfactual contribution *C*_*k*_ of program *k* is defined by

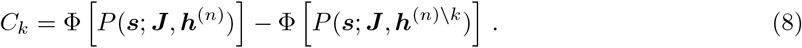

### Toggle switch model with time-dependent noises

We analyze the following toggle switch model with signal input and noise

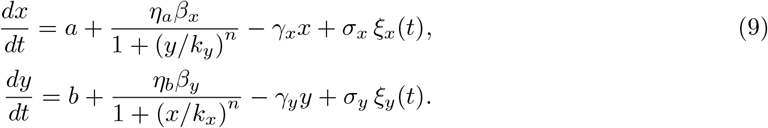

For simplicity, we assume *β*_*x*_ = *β*_*y*_ = *β, γ*_*x*_ = *γ*_*y*_ = *γ, k*_*x*_ = *k*_*y*_ = *k*. Noise amplitude 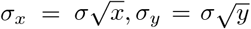 following the scaling of chemical reaction systems. *ξ*_*x*_, *ξ*_*y*_ are independent standard Wiener inputs. *a, b* ∈ [0, *β*] represent external signal activations. *η*_*a*_, *η*_*b*_ ∈ [0, 1] represent external signal inhibitions that reduce the production rate. The simulated model, unless otherwise noted, uses parameters *β* = 3, *k* = 1, *γ* = 1, *n* = 4, *σ* = 1.5. Simulations were performed by the Euler–Maruyama method with Δ*t* = 0.001. Noise decay in the annealing process follows the equation

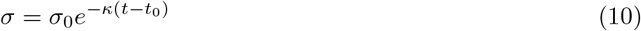

where *κ* = 0.2, and *t*_0_ = 5 to allow the system to arrive stationary. In the simulations for the signal input effect, signal ranges are chosen to cover the range of 10% 90% state X proportions. Specifically, we choose *a, b* ∈ [0, 0.5770] and *η*_*a*_, *η*_*b*_ ∈ [0.6185, 1].

To quantify the effective noise level, i.e., the rate at which the distribution approaches the stationary state, we analyze the transition rate *r* between the two attractor points. We first derive the dimensionless equation by taking 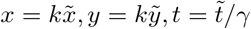, the equation now have the form

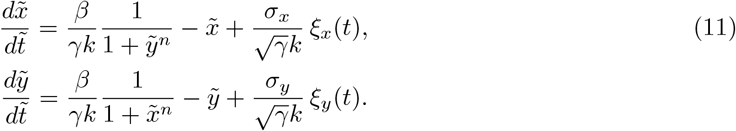

The final dimensionless equations have the form

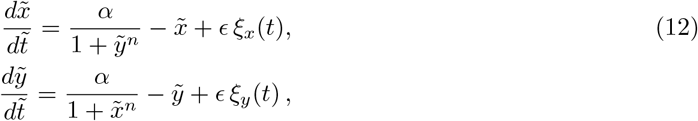

where 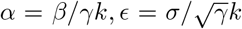. For simplicity, we treat the noise level as a constant. By the Freidlin-Wentzell quasipotential, the transition rate between the two attractor states asymptotically is *r* ≍ exp −Δ*S/ϵ*^2^, where Δ*S* is the height of the energy barrier of crossing the saddle point in the middle [6]. The toggle switch model does not follow a potential dynamics where *d****x****/dt* = _***x***_*U* + *σξ*_*t*_; therefore, the universal analytical form of the energy barrier height is not tractable. Nonetheless, we can analyze the proximity of the bifurcation point where the saddle and attractors are close enough to apply the Taylor expansion of the function.

By the symmetry of the equation, we always have the fixed point at *x* = *y* = *s* with *s* = *α/*(1 + *s*^*n*^). The Jacobian at the fixed point has the form

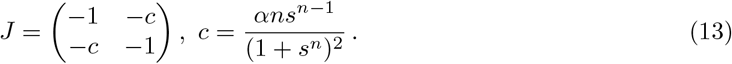

By changing *α*, the pitchfork bifurcation occur at *c* = 1 with the critical *α*_*c*_ = *n/*(*n* − 1)^1+1*/n*^. Then the dynamical system can be written in the normal form near the center manifold using the new coordinate

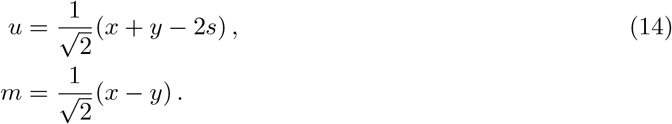

The attractor is always stable in the direction of *u*, and transitions from stable to unstable in the direction of *m* as *α* increases and passes *α*_*c*_. Near the bifurcation point with small *α* − *α*_*c*_, the derivatives of *m* follow the Landau theory of phase transition form (omitting noise for simplicity)

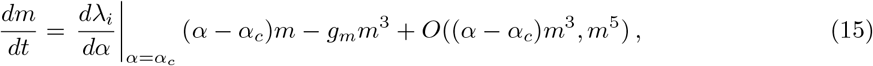

where *λ*_*i*_ = *c* − 1 is the eigenvalue of the unstable direction, *g*_*m*_ is a constant from the Taylor expansion of the dynamical system. The derivative corresponds to a double-well potential with the form

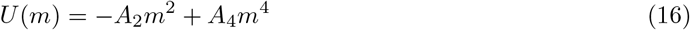

with energy barrier height scales with *A*^2^*/A*_4_. Therefore, we have Δ*S* ∝ (*α* − *α*_*c*_)^2^ and thus

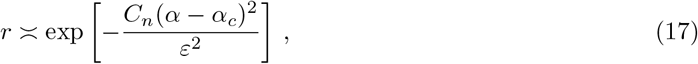

where *C*_*n*_ is a constant only depending on the Hill coefficient *n*. Inserting the original variables gives the final results of

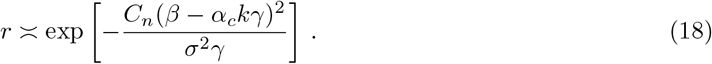

### Reconstruction of cell state trajectories with optimal transport

To compute the cell state transition vectors between cells in different experimental batches, we obtained the denoised, batch-corrected gene expression with scVI (scvi.model.SCVI.get normalized expression). We computed the optimal transport (OT) transition matrix between cells of adjacent time points (growth media day 0 to day 3 and day 3 to day 5; differentiation media day 0 to day 5) with the Python package wot for Waddington-OT with wot.ot.OTModel(epsilon=0.05, lambda1=1, lambda2=50, growth iters=3, local pca=10) [7]. In the transition matrix, each source cell could have nonzero transition fluxes to a group of similar target cells. We computed the flux-weighted average of target cell coordinates in the PCA space, and selected a single target cell that has the minimal distance to the target coordinate. Given a source cell PCA coordinate ***c***_source_ and its target ***c***_target_, we obtained one measurement of the vector field

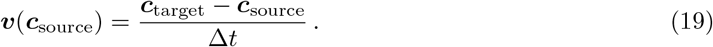

The computed vector field measurements of three dataset pairs were combined by local average, followed by local Gaussian smoothing with scipy.ndimage.gaussian filter(sigma=1.5).

### Quantification of state repulsion between fate regulators

One of the simplest models to quantify the mutual exclusivity of two genes is the two-node Ising model

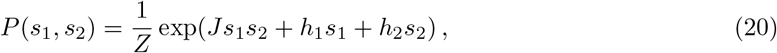

where *s*_1_, *s*_2_ ∈ {1, −1} are binary variables corresponding to on and off states of the two genes. *J* is the interaction between the two genes; *h*_1_, *h*_2_ are the activity level of the two genes; *Z* is the partition function. The interaction *J* can be directly solved from observed distributions by

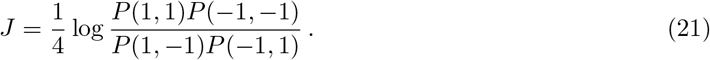

*J >* 0 indicates mutual activation and *J <* 0 indicates mutual inhibition. Therefore, we define the mutual repulsion index RI = −*J*. We use log_2_ base for better interpretability. The continuous gene expressions in the denoised gene expression matrix from scVI were partitioned into on and off binary states for each examined gene individually using 2-cluster KMeans clusterings (sklearn.cluster.KMeans).

### Analysis of the mouse cortex development dataset

We use the early development subset from a public mouse cortex development dataset [8], retaining unsorted cells from E11.5 to P0. We remove non-neuronal cell types and migrating neuron cells by excluding vascular, immune, GABAergic, and Cajal–Retzius populations, obtaining around 52,000 cells. Cell type annotations were inherited from the original study. Quality control and selection of 3,000 highly variable genes generally follow the procedures of our own dataset described above, with slight variation that (a) cell counts are normalized to the median library size (b) the gene detection threshold is 2% in any collection edge. We used the scVI architecture described above, with tissue region (‘roi’) as the batch variable (max epochs=100, batch size=128).

The gene-level D-SPIN network is constructed from the log-normalized expression following the same procedure described in our own datasets. We selected regulatory genes (TFs, kinases, and phosphatases) from databases with gene detection required in more than 2% of all cells [3, 4, 5]. Cell cycle and housekeeping genes are manually excluded from the network construction. The network of the remaining 325 genes was constructed with DSPIN.network inference(sample id key=‘Age’, method=‘pseudo likelihood’, directed=True, params=‘stepsz’: 0.04, ‘num epoch’: 1000).

For trajectory and regulator-pair analyses, we similarly used denoised, batch-corrected gene expression with scVI. The transition vector field is computed from optimal transport (ot.emd) between adjacent collection ages in the first 10 principal components of denoised expression. State repulsion between regulator pairs was computed similarly as described above. The thresholds of regulator expression were defined by the density minimum between the two highest expression peaks from kernel density estimates (‘scipy.stats.gaussian kde’). The dashed trends were obtained with Locally Weighted Scatterplot Smoothing (LOWESS) on equally spaced collection-age indices.

---

**Table 1.** Concentrations of signal ligands.

| Signal | Low | Medium | High |
| --- | --- | --- | --- |
| EGF/FGF2 | 0.8 ng/mL | 4 ng/mL | 20 ng/mL |
| BMP4 | 0 | 4 ng/mL | 20 ng/mL |
| Wnt (CHIR99021) | 0 | NA | 3 uM |
| PDGF | 0 | 4 ng/mL | 20 ng/mL |
| FGF9 | 0 | NA | 20 ng/mL |
| GM-CSF | 0 | NA | 3 ng/mL |
| IFNG | 0 | NA | 3 ng/mL |

**Table 2.** Information of single-cell profiling experiments.

| Sample name | Description | #Cell | Type |
| --- | --- | --- | --- |
| SVZ_D0_small | E18 subventricular zone | 563 | live |
| SVZ_D0 | E18 subventricular zone | 4419 | live |
| AdCo_small | Cortex from 8-week old mouse | 2541 | live |
| AdCo | Cortex from 8-week old mouse | 2511 | live |
| P4_1 | Whole brain from P4 mice | 3939 | live |
| P4_2 | Whole brain from P4 mice | 3562 | live |
| P4_3 | Whole brain from P4 mice | 4498 | live |
| SVZ_D3 | SVZ cells cultured 3 days in NPgrow media | 9862 | live |
| SVZ_D5 | SVZ cells cultured 5 days in NPgrow media | 1124 | live |
| SVZ_D7_small | SVZ cells cultured 7 days in NPgrow media | 512 | live |
| SVZ_D7 | SVZ cells cultured 7 days in NPgrow media | 3293 | live |
| SVZ_D18 | SVZ cells cultured 18 days in NPgrow media | 6235 | live |
| SVZ_D0_DIFF | SVZ cells directly cultured 5 days in NBco-culture media | 2680 | live |
| SVZ_D0_DIFF_small | SVZ cells directly cultured 5 days in NBco-culture media | 951 | live |
| NPC_D5_Growth | SVZ cells cultured 5 days in NPgrow media | 10470 | live |
| NPC_D5_Diff | 5 days in NPgrow media + 5 days in NBco-culture media | 3307 | live |
| NPC_MULT_1 | 5 days in NPgrow (minus EGF/FGF2) + signals (Batch1) | 8684 | fixed |
| NPC_MULT_2 | 5 days in NPgrow (minus EGF/FGF2) + signals (Batch2) | 11465 | fixed |

## Notes

### Competing Interest Statement

The authors have declared no competing interest.

### Summary of Updates

Tightened title and narrative; corrected typos.

